# Cyclic AMP competitively inhibits periplasmic phosphatases to coordinate nutritional growth with competence development of *Haemophilus influenzae*

**DOI:** 10.1101/2022.08.24.505078

**Authors:** Kristina Kronborg, Yong Everett Zhang

## Abstract

Natural competence is an important means of horizontal gene transfer that bacteria use to gain new physiological traits such as multiple drug resistance. Most naturally competent bacteria tightly regulate the window of competence state to maximize their ecological fitness under specific conditions. Here we study the inhibitory effect of purine nucleotides on the natural competence in *Haemophilus influenzae* Rd KW20. We first identified a periplasmic acid phosphatase AphA_Ec_ of *E. coli* as a new cyclic AMP (cAMP)-binding protein. cAMP competitively inhibits AphA_Ec_. Subsequently, we found that cAMP also competitively inhibits AphA_Hi_ and two additional periplasmic phosphatases NadN_Hi_ and Hel_Hi_ of KW20. Hel_Hi_ cleaves NADP to NAD, and NadN_Hi_ cleaves NAD to NMN and NR (nicotinamide ribose), providing the essential growth factor V for KW20. Consistently, we found that cAMP inhibits growth of KW20 in sBHI medium supplemented with NAD, but not NR. Moreover, the combined deletion of *aphA*_*Hi*_, *nadN*_*Hi*_, and *hel*_*Hi*_, but not the single or double deletion mutants, made KW20 immune to the inhibition of nucleotides on competence development. However, nucleosides still inhibited the competence of the triple mutant. Finally, cAMP in a dose-dependent manner restored the competence inhibited by the nucleotide GMP, but not by the nucleoside guanosine. Altogether, we revealed an antagonistic mechanism of cAMP and nucleotides in regulating cell growth and competence of *H. influenzae*. Similar mechanisms are discussed in other *H. influenzae* related organisms and *Vibrio cholerae*.

**Author summary:** *Haemophilus influenzae* is an important human pathogen, causing respiratory tract infection including pneumonia. Extensive drug resistance is observed in *Haemophilus influenzae* species, which is attributed to their well-described natural competence system. Natural competence is a physiological state that some bacteria, under certain conditions, become active to take up external DNA and integrate it into the chromosome. External DNA may contain an antibiotic resistance gene and thereby confer antibiotic resistance on *Haemophilus influenzae*. Therefore, it is important to understand how natural competence of *Haemophilus influenzae* is regulated by external cues. Previously, it was found that the secondary messenger cyclic AMP (cAMP) activates while nucleotides inhibit the competence development of *Haemophilus influenzae* Rd KW20. However, the interplay between cAMP and nucleotides is unclear. Here we show that cAMP competitively inhibits three periplasmic phosphatases of *Haemophilus influenzae* Rd KW20 and thereby inhibits the utilization of nutritional nucleotides and the essential growth factor NAD. Via this mechanism, cAMP activates the competence development of *Haemophilus influenzae* Rd KW20 only after external nucleotides are sufficiently depleted, coupling growth arrest with competence development. Similar mechanisms are anticipated to function in bacteria closely related to *Haemophilus influenza*e*e*, and also *Vibrio cholerae*, another important human pathogen.

## Introduction

In the phenomenon of natural competence, a bacterium initiates a genetic program to take up external DNA and integrate it into its chromosome. It is currently believed that natural competence is critical for the horizontal gene transfer, and thus bacterial genome evolution and the emergence of multidrug resistant (MDR) bacteria (1). Understanding when and how bacteria become competent is therefore essential to mitigate the detrimental effect of MDR bacteria in health care. Most of the naturally competent bacteria have a tightly regulated time window to become competent in response to environmental stresses, the regulatory mechanisms of which are intensively studied (2). Here, we studied the molecular mechanism of the inhibitory effect of nucleotides on the competence development in the model organism *Haemophilus influenzae* Rd KW20 (hereafter KW20) (3, 4).

The competence program of KW20 begins with the production of 3’,5’-cyclic AMP (cAMP) upon bacterial perception of stresses, such as exhaustion of carbon source at the end of the growth-phase in rich medium (5). In the laboratory, KW20 competence is often induced by shifting log-phase cells grown in rich medium to M-IV minimal medium that lacks carbon sources and consequently induces cAMP production (6). Additionally, high concentrations (1-10 mM) of exogenously added cAMP to growing cells induces competence of KW20 (7). cAMP binds to the carbon catabolite protein or cAMP receptor protein (CRP_Hi_), to stimulate the production of the master competence regulator Sxy_Hi_ (8, 9). Furthermore, the mRNA of *sxy*_*Hi*_ encodes a long non-coding 5’ sequence that is currently believed to perceive intracellular signals to control translation of *sxy*_*Hi*_ (9). Consistently, genetic changes in the *sxy*_*Hi*_ mRNA leader region produce constitutively competent KW20 mutants (5, 9). The ternary complex of cAMP-CRP_Hi_ -Sxy_Hi_ stimulates the expression of 25 genes that are involved in taking up external DNA and thus competence development in KW20 (10). Thus, cAMP and CRP_Hi_ link competence development to the quality of environmental nutrients in KW20.

In contrast to carbon starvation, the purine nucleotides AMP and GMP inhibit competence development of KW20 (4). Both purine nucleotides and the corresponding nucleosides inhibit competence development when they are added early during the M-IV medium induced competence program (3). Previous studies suggest a model wherein external purine nucleosides enter the cytosol of KW20 and participate in the purine nucleotide biosynthesis pathway. This metabolic change is perceived by factors including PurR, or the Sxy_Hi_ mRNA 5’-end structure to repress the translation of Sxy_Hi_ via a still mysterious mechanism that may involve a riboswitch (3). Nucleotides added in a later stage of M-IV induced competence program failed to inhibit competence and the constitutively active Sxy_Hi_ mutants are not inhibited by nucleotides (3), indicating that nucleosides affect Sxy_Hi_ production. However, the connections between cAMP, nucleotide metabolism, and competence development are incompletely understood.

Besides competence, KW20 has several unique features tied with the metabolisms of nucleotides and other metabolites with similar chemical structures. Firstly, KW20 encodes an incomplete pathway to synthesize NAD (nicotinamide adenine dinucleotide), an essential molecule for all organisms. Therefore, NAD (the so-called factor V for KW20), NMN (nicotinamide mononucleotide), or NR (nicotinamide riboside) is required to support the growth of KW20. A periplasmic phosphatase NadN_Hi_ is essential for the conversion of NAD to NMN and subsequently to NR (11), which then traverses into the cytosol via PnuC (11, 12). Secondly, the oxygen carrier hemin (essential growth factor X) is required for KW20 to grow. The utilization of hemin was thought to require a lipoprotein Hel_Hi_ or e(P4). However, it was later found that Hel_Hi_ converts NADP to NAD, and potentially also further to NMN and NR (13, 14). In addition, both NadN_Hi_ and Hel_Hi_ show phosphatase activities towards nucleotides (13, 14). Finally, KW20 is known to require external pyrimidines to grow. In total, nucleotide metabolism affects both cell growth and competence development in KW20, but the detailed underlying mechanisms remain to be clarified.

Here, we started with a systematic screening of cAMP-binding proteins in *Escherichia coli* and, surprisingly, identified the periplasmic acid phosphatase AphA_Ec_. We show that cAMP competitively inhibits the AphA_Ec_ catalytic activity. Furthermore, we find that cAMP competitively inhibits AphA_Hi_, and in addition, also Hel_Hi_, and NadN_Hi_. We find that a combined deletion of the three genes *aphA*_*Hi*_, *nadN*_*Hi*_, *hel*_*Hi*_ make KW20 immune to the inhibition of nucleotides on competence development; however, nucleosides still inhibited competence of the triple mutant. Finally, we find that cAMP restores the competence inhibited by nucleotides, but not by nucleosides. Altogether, we reveal and discuss the role of cAMP and the three phosphatases in coupling cell growth with competence development in KW20 and other related organisms.

## Results

### A proteome-wide screening identified the periplasmic acid phosphatase AphA as a novel cAMP binding protein in *E. coli*

The well-studied second messenger cAMP is critical for bacterial carbon metabolism, pathogenesis, and virulence (15). Presently, the only known target protein of cAMP in *E. coli* is CRP_Ec_ (or CAP). To explore if cAMP has additional effector proteins, we performed a proteome-wide screening of cAMP binding proteins using DRaCALA (16, 17). First, cAMP was synthesized from p^32^-α-ATP by using the truncated recombinant *E. coli* CyaA protein (18) (conversion ratio > 85%, Figure S1A). The ordered ASKA strain collection (19) was used to overexpress proteins of *E. coli* K-12 MG1655, and whole cell lysates were prepared as before (20). Radioactively labelled p^32^-α-cAMP was then mixed with individual whole lysates to screen for cAMP binding proteins. CRP_Ec_ gave a strong binding signal (Figure 1A), validating the screening method. Besides CRP_Ec_, AphA_Ec_, a non-specific periplasmic phosphatase, also gave a strong binding signal (Figure 1B; see other DRaCALA screening plates in Supplementary document S1).

**Figure 1.**
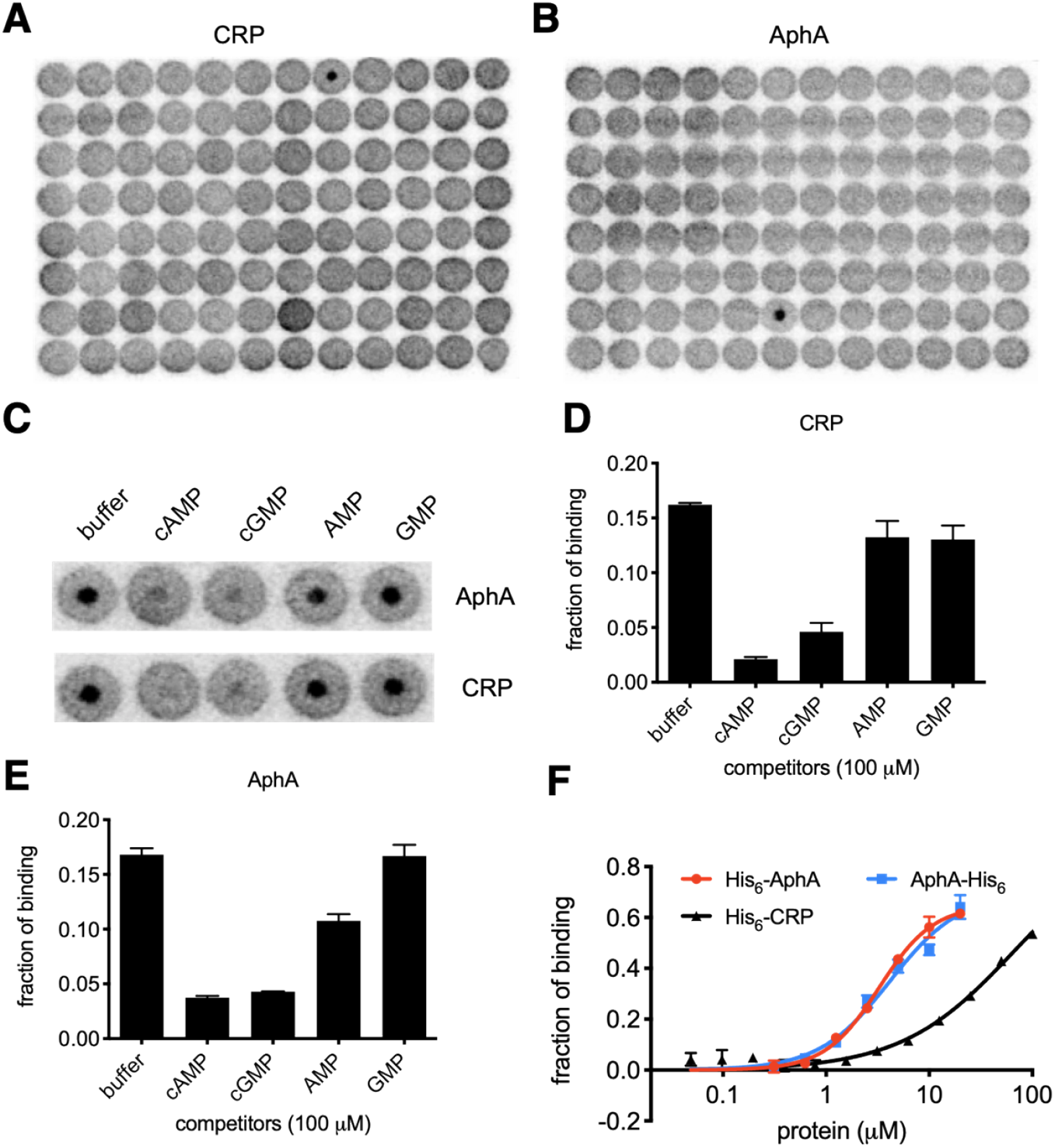
cAMP binds to AphA of *E. coli*. (**A, B**) Autoradiography of the two DRaCALA screening plates that identified CRP_Ec_ and AphA_Ec_ as cAMP binding proteins. (**C**) DRaCALA-based competition assay by using whole cell lysates harboring overproduced AphA_Ec_ or CRP_Ec_, with the presence of buffer, cold cAMP, cGMP, AMP and GMP (each at 100 µM). (**D, E**) Quantitation of the p^32^-α-cAMP binding fractions from panel (C). Two biological replicates were included. The average of binding fraction and the standard deviation of mean are shown. (**F**) Determination of the K_d_ values of cAMP binding to His_6_-AphA, AphA-His_6_, and His_6_-CRP proteins, by using DRaCALA (16, 17). Two biological replicates were performed, and the average of binding fraction and the standard deviation of mean are plotted.

To confirm the binding, the two ASKA strains overproducing CRP_Ec_ or AphA_Ec_ were used in a DRaCALA-based competitive binding assay. Various cold nucleotides (100 µM) were added to the binding reactions. Figure 1C and 1D,1E show that cold cAMP, and also cyclic GMP (cGMP), effectively outcompeted the binding of p^32^-α-cAMP to both CRP_Ec_ and AphA_Ec_. On the other hand, the known AphA_Ec_ substrates AMP and GMP were less effective (see below), indicating that the binding of cAMP to CRP_Ec_ and AphA_Ec_ was specific.

### cAMP binds to AphA_Ec_ with high affinity

AphA_Ec_ is a periplasmic protein with an N-terminus (Nt) signal peptide. The ASKA plasmid produces a recombinant AphA_Ec_ with a six-histidine tag N-terminal to the signal peptide and a C-terminal (Ct) GLCGR peptide (19). To understand how cAMP binds to AphA_Ec_, we first removed the signal peptide and fused a six-histidine (His_6_) tag at either the Ct-or Nt of AphA_Ec_ (AphA_Ec_-His_6_ and His_6_-AphA_Ec_, respectively). Both proteins were purified to homogeneity via tandem affinity purification and size exclusion chromatography (SEC). The SEC profiles (Figure S1B) suggested that AphA_Ec_-His_6_ formed a dimer whereas His_6_-AphA_Ec_ formed a tetramer. Several crystal structures of AphA_Ec_ homologs were reported (21–23) and inspection of a published crystal structure of AphA_Ec_ homolog (PDB 2B82) suggested that a Ct His_6_ tag potentially leads to a steric clash between the neighboring two monomers (Figure S1D, S1E), destabilizing the tetrameric configuration. Both *E. coli* protein variants were therefore purified (Figure S1C) and used to measure the binding affinity of cAMP using DRaCALA (Figure 1F). Low micromolar range K_d_ values were obtained for both proteins (4.4 ± 0.4 µM and 3.3 ± 0.4 µM for AphA_Ec_-His_6_ and His_6_-AphA_Ec_, respectively), suggesting a high binding affinity of cAMP to AphA_Ec_ and that the His_6_ tags and multimeric states did not affect the binding of cAMP to AphA_Ec_ dramatically. As a control, the His_6_ tagged CRP_Ec_ from the ASKA library binds to cAMP with a K_d_ value of 53 ± 20 µM (Figure 1F), similar to previously reported values (24, 25). cAMP is chemically similar to AMP, a substrate of AphA_Ec_. We found, however, that AphA_Ec_ did not cleave cAMP (Figure S1F), consistent with previous studies (26). We then performed the DRaCALA competitive binding assay again with the purified AphA_Ec_ proteins and found that 100 µM of cAMP and cGMP, but not the substrates AMP, GMP or other purine nucleotide di-, tri-phosphates, were able to outcompete the binding of p^32^-α-cAMP by CRP_Ec_ and AphA_Ec_ (Figure S1G-I). Since low micromolar range K_m_ values (3 and 15 µM, (26)) of AMP and GMP were reported for AphA_Ec_, the data appear to indicate that cAMP binds to a site different from the catalytic pocket of AphA_Ec_ (see below).

### cAMP competitively inhibits the acid phosphatase activity of AphA_Ec_

As a non-specific acid phosphatase, AphA_Ec_ degrades many nucleotide monophosphates to generate nucleoside and orthophosphate (26). To understand the effect of cAMP on the AphA_Ec_ activity, we performed a phosphatase assay of AphA_Ec_ by using pNPP (p-Nitrophenyl Phosphate) as the substrate (26). Cleavage of pNPP releases a phosphate and pNP, a yellow chemical with a maximal absorption at 405 nm, which could be used to quantitate the reaction (26). His_6_-AphA_Ec_ was firstly tested at the reported optimal pH 5.6 and cAMP (100 µM) inhibited the catalytic activity of His_6_-AphA_Ec_ two-fold. A similar assay was performed at a higher pH 8.0 and a ten-fold inhibition was observed despite the lower activity of His_6_-AphA_Ec_ (Figure S2A). However, we found that cAMP stimulated the catalytic activity of AphA_Ec_-His_6_ at pH 5.6 in a dose dependent manner (Figure S2B). Despite the low fold of stimulation, the effect was highly reproducible (Figures S2C, S2D). Of note, the activity of AphA_Ec_-His_6_ was lower than His_6_-AphA_Ec_, and cGMP stimulated AphA_Ec_-His_6_ activity as well (Figure S2B). Despite this observation, the dimeric configuration of AphA_Ec_-His_6_ is likely a non-natural state (see below) and this artificial phenomenon was not studied further.

Given the opposite effects of cAMP on the Nt and Ct His_6_ tagged AphA_Ec_, we constructed a tag-less AphA_Ec_ to clarify the regulatory effect of cAMP. To do this, we cloned AphA_Ec_ with a Nt His_6_-SUMO tag, which was cleaved off by using the SUMO specific protease Ulp1 (His_6_-Ulp1). The SDS-PAGE gel (Figure S3A) showed that the His_6_-SUMO tag was successfully cleaved off to generate a protein around 25 kDa, matching the expected tag-less AphA_Ec_. The SEC profile revealed a peak with the calculated size of 100 kDa, a tetrameric form of tag-less AphA_Ec_ (Figure S3B). These data confirm that the tetrameric form is the natural state of AphA_Ec_. Subsequently, we performed the pNPP phosphatase assay by using the tag-less AphA_Ec_ and varied concentration of cAMP (Figure 2A,2B). The Michaelis-Menten curves were fitted with different models of inhibition. An allosteric sigmoidal fit was found to be the best, with the fitted Hill coefficient between 1 – 1.3. Moreover, fitting to the different models of inhibition and a Lineweaver–Burk plot (Figure 2B) indicated that cAMP probably inhibits the catalytic activity of AphA_Ec_ in a competitive manner (K_i_ = 3.9 ± 0.3 µM). We then performed a kinetic study of the tetrameric His_6_-AphA_Ec_ and obtained very similar curves and K_i_ value (11 ± 0.6 µM) (Figure S3C, S3D). These data suggest that the Nt histidine tag does not greatly affect the activity of cAMP on AphA_Ec_. We therefore used the His_6_-AphA_Ec_ for the subsequent experiments.

**Figure 2.**
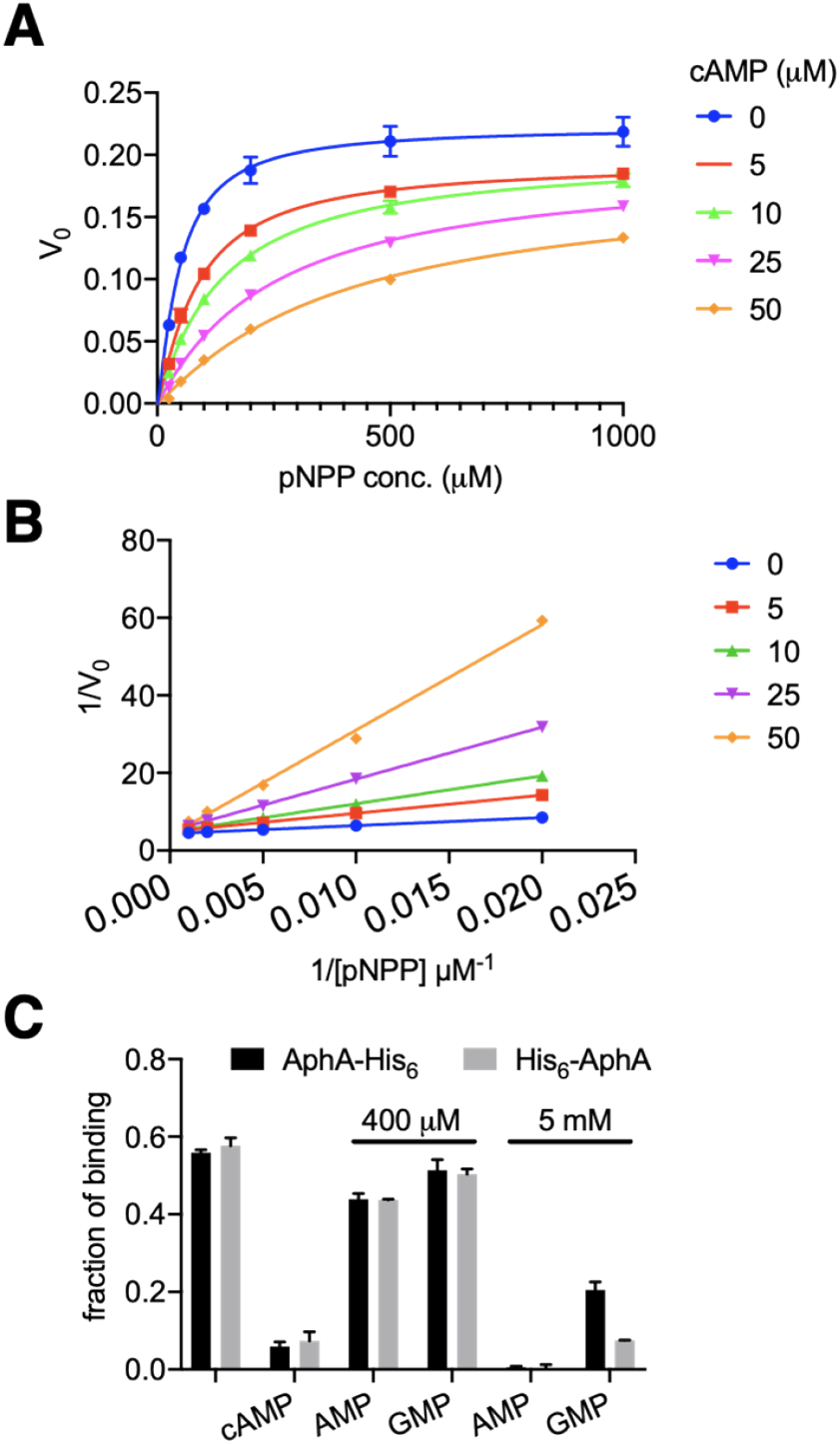
cAMP competitively inhibits the phosphatase activity of AphA_Ec_ of *E. coli*. (**A**) Michaelis-Menten curves (in three biological replicates) and (**B**) the Lineweaver–Burk plot of tag-less AphA_Ec_ cleaving pNPP (p-Nitrophenyl Phosphate) in the absence or presence of various concentrations of cAMP. (see text and Method for details). (**C**) Quantitation of DRaCALA competition assay by using purified His_6_-AphA_Ec_ and AphA_Ec_-His_6_ proteins, without or with cold cAMP (100 µM), AMP (400 µM, 5 mM) and GMP (400 µM, 5 mM). Two biological replicates were performed, and the average and standard deviation of mean are shown.

The competitive inhibition of AphA_Ec_ activity contradicts the DRaCALA competitive binding assay (Figure 1C-E), which indicates an allosteric binding of cAMP. To study this further, we performed DRaCALA competitive binding assay by using even higher (400 µM) concentrations of AMP and GMP and observed a slight reduction of the binding fractions with AMP, but not GMP (Figure 2C). We then used 5 mM of AMP and GMP. This time, the binding fractions dropped to nearly zero for AMP and to the similar level of 100 µM cAMP for GMP (Figure 2C), indicating that AMP has a higher affinity to AphA_Ec_ than GMP as reported (26).

A further search of AphA_Ec_ homologs in the PDB database found the AphA from *Salmonella typhimurium* (AphA_St_, 1Z5U), which was serendipitously crystalized in complex with cAMP (Figure S3E, S3F). AphA_St_ shares extensive amino acid sequence similarity (89% amino acid sequence identity) and striking structural similarity with AphA_Ec_ (2B82, Figure S3E, S3F, rmsd = 0.331). Superposition of both protein structures indicates that cAMP can bind to the catalytic site of AphA_Ec_. Altogether, these data suggests that cAMP binds to the active site of AphA_Ec_ and strongly inhibits its phosphatase activity.

### AphA from *H. influenza* Rd KW20 is competitively inhibited by cAMP

The authentic physiological function of AphA_Ec_ seems to cleave nucleotides and provide them as carbon source for *E. coli* (27), despite the report that AphA_Ec_ may bind to hemi-methylated DNA in *E. coli (28)*. On the other hand, AphA homolog from *Salmonella typhimurium* (AphA_St_) was reported to facilitate the uptake of NAD (29). Moreover, several pieces of evidence suggest that AphA in *Hemophilus influenzae* (AphA_Hi_) is functionally related to the natural competence in *H. influenzae*. 1) Nucleotides and nucleosides inhibit the natural competence of *H. influenzae* (4); 2) The promoter region of *aphA*_*Hi*_ is predicted to encode a binding site of PurR_Hi_, which regulates natural competence in *H. influenzae* (3). 3) Two positively charged surface areas exist and are conserved on AphA homolog proteins (Figure S4A,B, circulated). Therefore, we turned to study the potential function of AphA_Hi_ in natural competence of KW20.

Similar as the AphA_St_ (PDB, 1Z5U), AphA_Hi_ is a close homolog of AphA_Ec_ with 49% identity and 69% similarity at the primary sequence level. AphA_Hi_ is thus anticipated to have a similar structural fold, raising the possibility that cAMP also inhibits AphA_Hi_. To test this, we firstly purified the His_6_-AphA_Hi_ and tested its binding to cAMP via DRaCALA. His_6_-AphA_Hi_ binds to cAMP with a high affinity (K_d_ = 1.03 ± 0.04 µM) (Figure 3A). We then found that cAMP inhibits His_6_-AphA_Hi_ at both pH 5.6 and 8.0 (Figure 3B). Further enzyme kinetic analysis showed that cAMP inhibits His_6_-AphA_Hi_ in a competitive manner (K_i_ = 6.9 ± 0.7 µM) (Figure 3C,3D). This shows that cAMP strongly inhibits AphA_Hi_.

**Figure 3.**
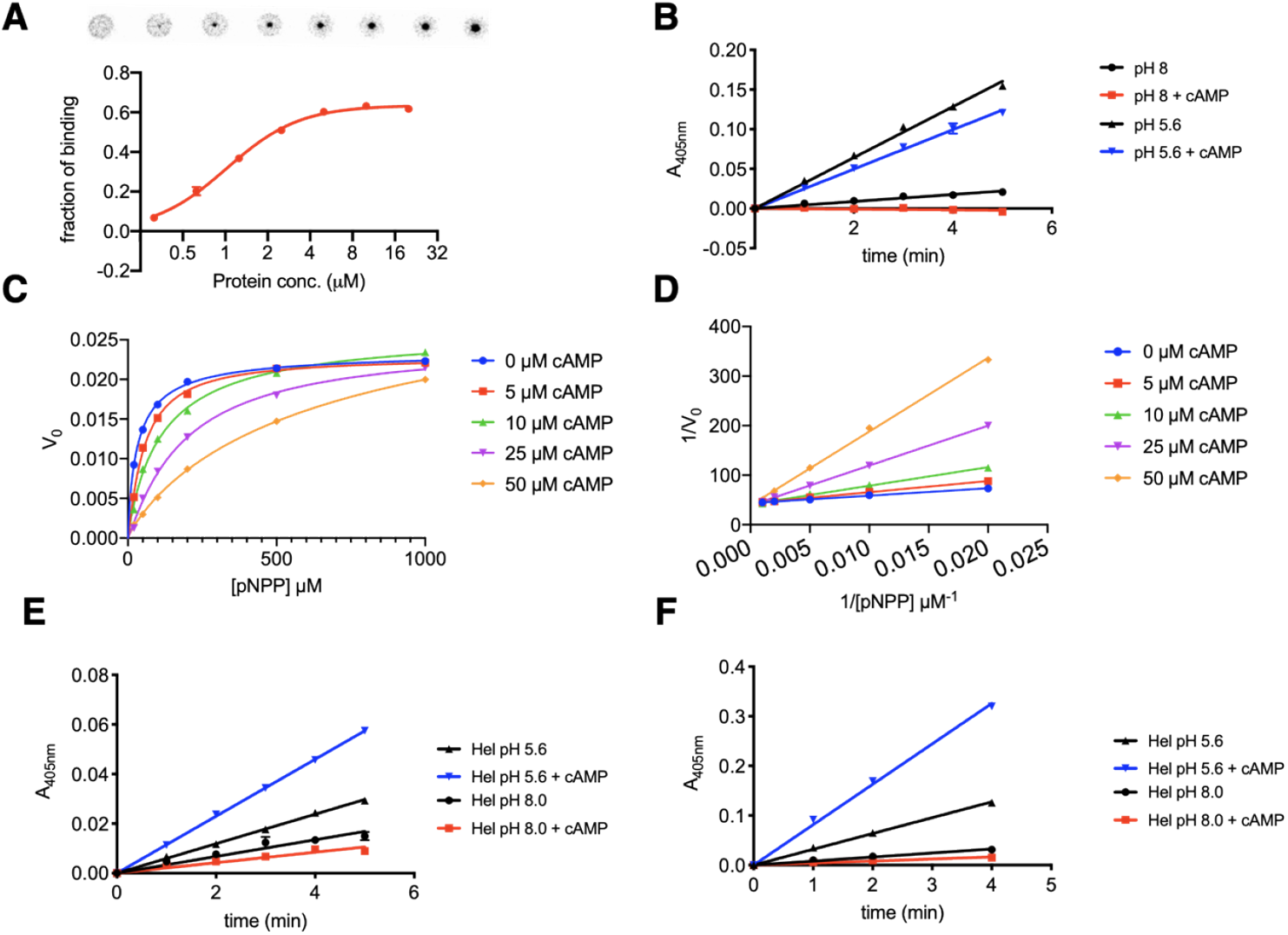
cAMP strongly binds and competitively inhibits the AphA_Hi_ from *H. influenzae*. (**A**) DRaCALA assay to measure the binding affinity of His_6_-AphA_Hi_ to ^32^p-cAMP. Two biological replicates were performed, and the average and standard deviation of mean are shown. (**B**) The acid phosphatase assay was performed by using pNPP (p-Nitrophenyl Phosphate, 1 mM) as substrate, 15 nM of His_6_-AphA_Hi_ protein at pH 8.0 and 5.6, without or with 100 µM cAMP (see text and Method for details). (**C**) Michaelis-Menten curves and (**D**) the Lineweaver–Burk plot of His_6_-AphA_Hi_ cleaving pNPP in the absence or presence of varied concentrations of cAMP. (**E, F**) The acid phosphatase assay of the His_6_-Hel_Hi_ protein (**E**,15 nM) and a tag-less Hel_Hi_ protein (**F**,15 nM) was performed by using pNPP (p-Nitrophenyl Phosphate, 1 mM) as substrate, at pH 8.0 and 5.6, without or with 100 µM cAMP (see text and Method for details). At least two biological replicates were performed and the average and the standard deviation of mean are plotted.

### His_6_-AphA_Hi_ does not bind to double-or single-strand DNA

The periplasmic AphA_Hi_ may bind directly to imported DNA molecules. To test this, we amplified a 200-bp long DNA sequence from KW20, which contains an KW20 specific uptake-sequence (UPS) ACCGCACTT. Gel retardation assay showed that His_6_-AphA_Hi_ (up to 50 µM) did not shift the DNA band, regardless the presence of cAMP (Figure S4C). We then denatured the dsDNA to ssDNA (see method for preparation) and tested the binding again. As a control, we used the recombinant *E. coli* single-strand DNA binding protein (SSB, P0AGE0). His_6_-SSB_Ec_ shifted the ssDNA from as low as 0.13 µM (Figure S4D). However, we still did not see a shifted band with His_6_-AphA_Hi_ (up to 50 µM). These data suggest that His_6_-AphA_Hi_ does not directly bind to dsDNA or ssDNA.

### Δ*aphA*_*Hi*_ is not defective in starvation medium M-IV induced competence

To test directly if AphA_Hi_ is involved in KW20 competence, we constructed the *aphA*_*Hi*_ deletion strain Δ*aphA*_*Hi*_*::cat*, where *aphA*_*Hi*_ was replaced with a chloramphenicol resistance marker. Δ*aphA*_*Hi*_*::cat* grows similar as the WT KW20 in sBHI rich medium (Figure S5A). We then tested the competence phenotype under several conditions, i.e., during growth into stationary phase in sBHI medium, M-IV starvation medium induced competence, and cAMP (1 mM) induced competence of Log phase sBHI culture. Nucleotides (and nucleosides) inhibit the natural competence in *H. influenzae* (4). Given the fact that AphA degrades nucleotides, we also tested the role of AphA_Hi_ in AMP mediated inhibition of KW20 competence. However, there was no obvious difference in competence efficiency between wt KW20 and Δ*aphA*_*Hi*_*::cat* strains under all the tested competence conditions (data not shown).

### cAMP affects the catalytic activity of Hel_Hi_ in a pH-dependent manner

The current model of nucleotide mediated inhibition of KW20 natural competence propose that extracellular nucleotides are degraded to nucleosides, which enter the cytosol to inhibit the translation of the master regulator of competence, Sxy_Hi_ (3). Besides AphA_Hi_, KW20 encodes two additional phosphatases, the periplasmic NadN (NadN_Hi_) and the outer membrane anchored Hel (Hel_Hi_, e(P4)). Hel_Hi_ is a close structural homolog of AphA (30), and dephosphorylates NADP, NMN and nucleotides (31). NadN_Hi_ first degrades NAD to NMN and AMP, then both NMN and AMP are further dephosphorylated by NadN_Hi_ into nicotinamide riboside (NR) and adenosine, which traverse the inner membrane (11). Therefore, NadN_Hi_ and Hel_Hi_ are functionally redundant with AphA_Hi_ regarding dephosphorylation of the compounds mentioned and thereby potentially the inhibition of competence as well.

To test if cAMP also competitively inhibits NadN_Hi_ and Hel_Hi_, we purified His_6_-NadN_Hi_ and His_6_-Hel_Hi,_ and performed phosphatase assay with pNPP and cAMP. Surprisingly, cAMP inhibited the phosphatase activity of His_6_-Hel_Hi_ at pH 8.0, but stimulated it at pH 5.6 (Figure 3E). To rule out potential effect of the histidine tag, we purified a tag-less Hel_Hi_ (Materials & Methods). However, the pH dependent effect of cAMP still holds for the tag-less Hel_Hi_ (Figure 3F). The physiological niche of KW20, mucin, has a pH range of 6-7, and it thus remains plausible that cAMP inhibits the phosphatase activity of His_6_-Hel_Hi_ under its physiological conditions.

### cAMP competitively reduces the growth-rate of KW20 in sBHI supplemented with NAD

The recombinant His_6_-NadN_Hi_ proteins purified from *E. coli* BL21 DE3 did not show any activity toward pNPP on our hands (data not shown). We thus turned to a different approach. The NAD degrading activity of NadN_Hi_ is essential for KW20 growth in sBHI (13) because KW20 cannot synthesize NAD. We found that when *hel*_*Hi*_ was deleted from either wt KW20 or Δ*aphA*_*Hi*_*::Cam* strain, both Δ*hel*_*Hi*_ and Δ*hel*_*Hi*_ Δ*aphA*_*Hi*_ strains were able to grow on sBHI medium supplemented with NAD. However, when *nadN*_*Hi*_ was further deleted, NAD did not support cell growth anymore; instead, NR was required for the growth of Δ*hel*_*Hi*_Δ*nadN*_*Hi*_ and Δ*hel*_*Hi*_Δ*nadN*_*Hi*_Δ*aphA*_*Hi*_. This confirmed the previously reported key function of NadN_Hi_ to convert NAD to NR (11), which enters the cytosol to support KW20 growth. We took advantage of this phenomenon to test if cAMP inhibits NadN_Hi_ function. In this scenario, the KW20 cells, by analogy, are viewed as a bag of enzymes functioning to utilize the essential factor V NAD (i.e., the substrate, in µM) for supporting cell growth (i.e. the reaction product of KW20 growth-rate, in cell division times per hour) and NadN_Hi_ is the first and rate-limiting enzyme. If cAMP inhibits the NAD-cleaving activity of NadN_Hi_, less NMN and NR will be produced, thus reducing the growth-rate of KW20. In this view, the growth-rate is proportional to the activity of NadN_Hi_.

Since high concentration of cAMP (1-10 mM) induces competence and inhibits the growth of KW20, we tested a much lower range of cAMP (from 1-500 µM). We measured the growth-rate of KW20 in sBHI supplemented with varied concentrations of the essential growth factors, i.e., the ‘substrates’ NAD, NR and hemin, in the presence of different levels of cAMP (Figure 4A-4C, S6). To analyze if cAMP competitively inhibits cell growth-rate and thus NadN_Hi_ activity, a double-reciprocal Lineweaver–Burk plot of the doubling time (i.e., the reciprocal of growth-rate, in hour per cell division) and substrate concentrations (i.e., in 1/[substrate concentration, µM]) was performed. Figure 4D showed characteristic straight lines converging to the Y-axis and indicates that cAMP reduces the growth-rate in a competitive manner when NAD was studied. Conversely, when NAD was replaced with NR or when the other essential growth factor hemin was assayed, the plotted curves were not consistent with a competitive model (Figure 4E, 4F). These data suggest that cAMP competitively inhibits the function of NadN_Hi_ in the cleavage of NAD to NMN, and/or NMN to NR.

**Figure 4.**
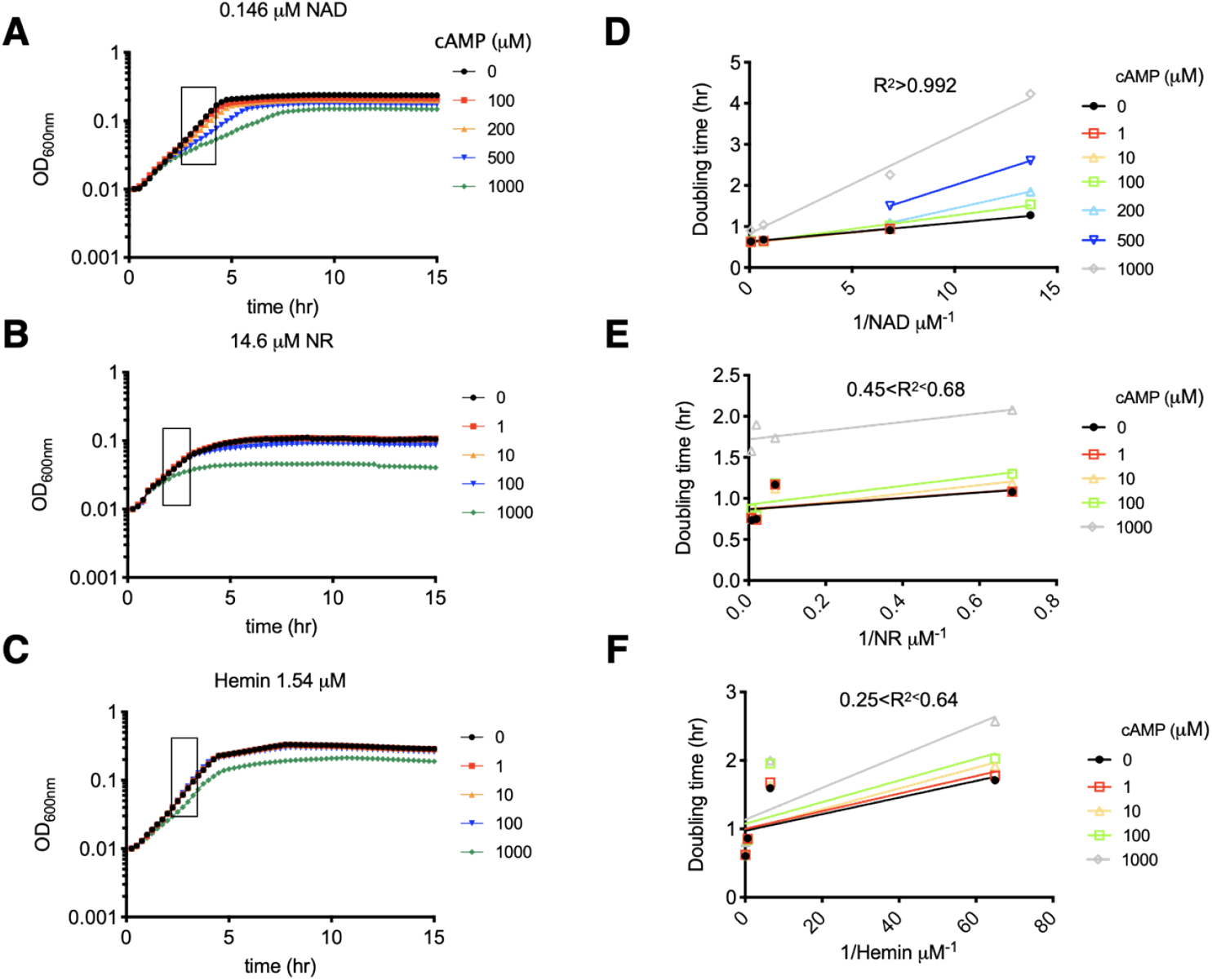
cAMP competitively inhibits *H. influenzae* Rd KW20 growth-rate in sBHI supplemented with NAD. (**A-C**) Representative growth curves of KW20 in sBHI medium supplemented with varied concentrations of the essential growth factors, i.e., NAD (**A**, 0.146 µM), NR (**B**, 14.6 µM) and hemin (**C**, 1.54 µM) in the presence of different concentrations of cAMP (µM). The framed regions of exponential phase growth data were used to calculate the growth-rates. See all growth curves in supplementary Figure S6. Three biological replicates were performed. (**D-F**) The double-reciprocal Lineweaver–Burk plot of the doubling time (i.e., the reciprocal of growth-rate, in hour) and the essential growth factors (in 1/[concentration of NAD/NR/hemin]) of wt *H. influenzae* were plotted. The R^2^ values of the linear fitting results are annotated above each diagram. The average growth-rate out of the three biological replicates (A-C) was used to calculate the doubling times.

### The triple mutant ΔnadN_Hi_Δhel_Hi_ΔaphA_Hi_ is refractory to the inhibitory effect of nucleotide on M-IV induced competence

The above data showed that cAMP inhibits the phosphatase activities of AphA_Hi_, NadN_Hi_ and Hel_Hi_. Given the redundant activities, we propose that AphA_Hi_, NadN_Hi_, and Hel_Hi_ altogether degrade nucleotides to nucleosides, to inhibit natural competence in KW20. We first found that the growth dynamics are similar in sBHI supplemented with NR (sBHI+NR) for wt, Δ*hel*_*Hi*_Δ*nadN*_*Hi*_ and Δ*hel*_*Hi*_Δ*nadN*_*Hi*_Δ*aphA*_*Hi*_ mutants, confirming that the main function of NadN_Hi_ is for utilizing NAD (Figure S5B). The three strains were grown in sBHI+NR broth to Log phase and shifted to M-IV starvation medium to induce competence. Fifteen minutes after the shift, various concentrations of AMP were added. As shown in Figure 5, AMP from as low as 10 µM reduced the M-IV induced competence by two-orders of magnitude in wt KW20. The triple mutant was completely refractory to the inhibitory effect of AMP (up to 4 mM tested), and the Δ*hel*_*Hi*_Δ*nadN*_*Hi*_ double mutant showed an intermediate effect (Figure 5A). These data suggest that AphA_Hi_, NadN_Hi_ and Hel_Hi_ all contribute to the inhibitory effect of AMP on KW20 competence. By contrast, M-IV induced competence of both the double and triple mutants were completely inhibited by adenosine (from 10 µM, Figure 5B).

**Figure 5.**
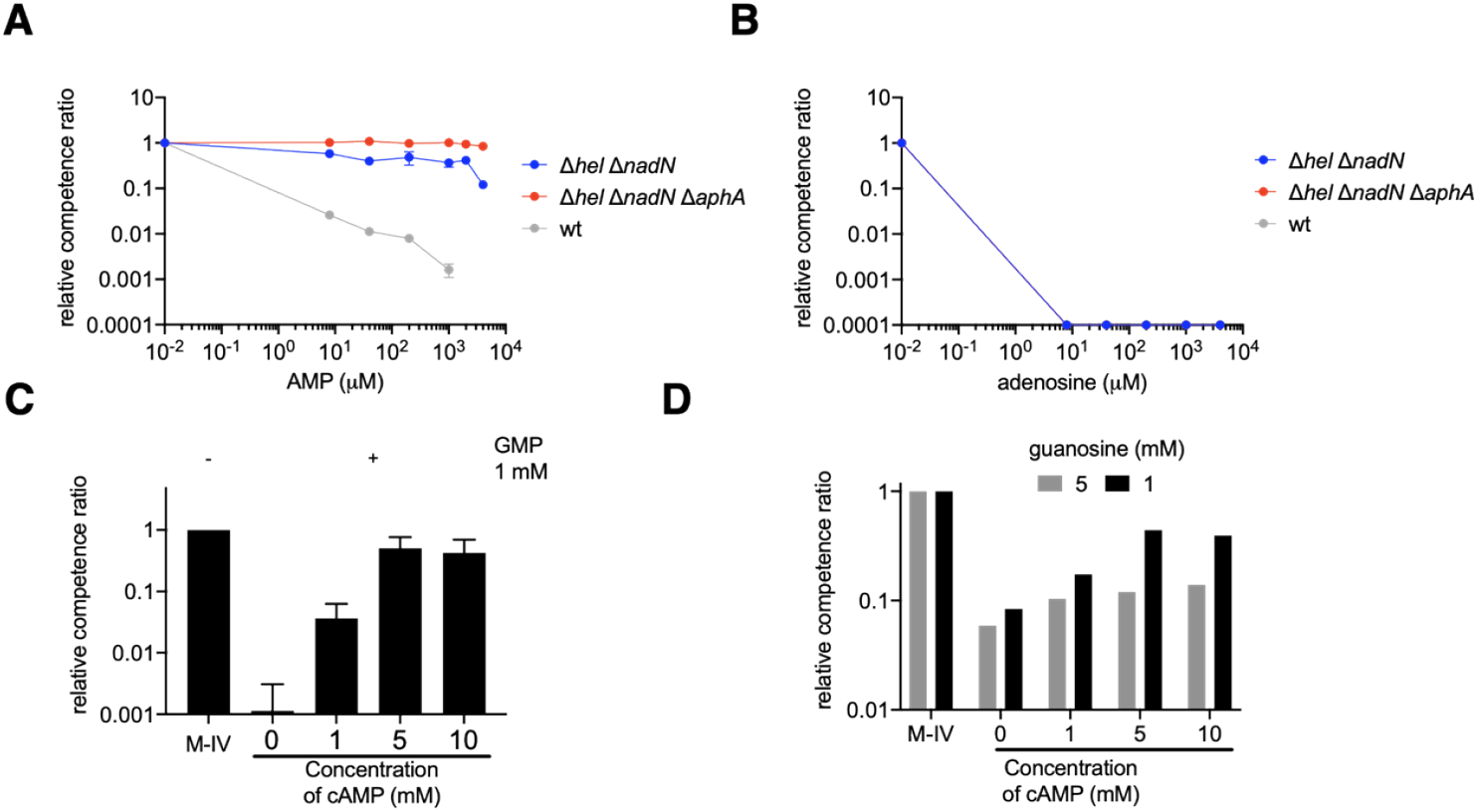
The M-IV induced competence development in *H. influenzae* Rd KW20 is affected by cAMP, nucleotides, nucleosides and AphA_Hi_, NadN_Hi_, Hel_Hi_. (**A**) Relevant competence ratios of wt, Δ*hel*_*Hi*_Δ*nadN*_*Hi*_, Δ*hel*_*Hi*_Δ*nadN*_*Hi*_Δ*aphA*_*Hi*_ in the presence of varied AMP concentration. The competence level of wt KW20 in the absence of AMP was used to normalize the competence of other strains. An arbitrary value, 0.01 µM, of AMP was used to plot the data obtained when AMP was absent. (**B**) Similar as panel A, except that the competence was determined in the presence of varied concentrations of adenosine. Two biological replicates were performed, and the average and standard deviation of mean are shown. (**C**) Relevant competence ratios of wt KW20 in the absence and presence of 1 mM GMP, and varied concentrations of cAMP. Three biological replicates were performed, and the average and standard deviation of mean are shown. (**D**) Similar as panel C, except that guanosine at 1 or 5 mM was used to inhibit KW20 competence.

Since cAMP competitively inhibits the activities of AphA_Hi_ and NadN_Hi_, we added cAMP (1-10 mM) in the assay to see if cAMP counteracts AMP in KW20 competence development. We observed that the AMP inhibited competence of wt KW20 was not restored by cAMP, consistent with previous report (4) (data not shown; see discussion below). Instead, we tested if cAMP restores the competence inhibited by GMP (1 mM), which has a weaker ability than AMP to outcompete the binding of ^32^p-cAMP to AphA (Figure 2C) and to inhibit competence (4). We found that cAMP restored the inhibited competence in a dose dependent manner (Figure 5C), consistent with previous report (4). Finally, we argue that nucleosides should still inhibit KW20 competence despite the presence of cAMP. Indeed, cAMP at the highest concentration (10 mM) only partially restored the KW20 competence inhibited by 1 mM guanosine, and the restoration was worse with 5 mM guanosine (Figure 5D). Taken together, these data demonstrate the roles of AphA_Hi_, NadN_Hi_ and Hel_Hi_ in the utilization of nucleotides and regulation of *H. influenzae* KW20 competence.

## Discussion

Development of bacterial competence is often characterized by a simultaneously inhibited cell growth (2), which results from an exhaustion of a key nutrient, typically a carbon source that consequently stimulates the production of cAMP. In *H. influenzae* KW20, competence is also inhibited by nucleotides, especially AMP and GMP, and their corresponding nucleosides, but not the nucleobases (4). KW20 is a fastidious bacterium requiring several essential factors, including NAD, hemin, and pyrimidines, to grow. How KW20 perceives and integrates these nutritional signals with competence development remains incompletely understood. In this study, we found via *in vitro* and *in vivo* methods that cAMP competitively inhibits the periplasmic non-specific phosphatases AphA_Hi_ and Hel_HI_ in KW20. Importantly, we showed that cAMP competitively inhibits the KW20 growth-rate in sBHI medium supplemented with NAD, but not NR, strongly suggesting that cAMP inhibits NadN_Hi_ as well. Since NadN_Hi_ also degrades a variety of nucleotides, it is imaginable that cAMP binds to the active site and competitively inhibits its activity. Moreover, only the triple deletion mutant Δ*hel*_*Hi*_Δ*nadN*_*Hi*_Δ*aphA*_*Hi*_ was refractive to the inhibitory effect of AMP on M-IV induced competence of KW20, consistent with the redundant activities of the three periplasmic phosphatases. We therefore propose a model of cAMP and the three phosphatases in coordinating KW20 cell growth and competence development (Figure 6).

**Figure 6.**
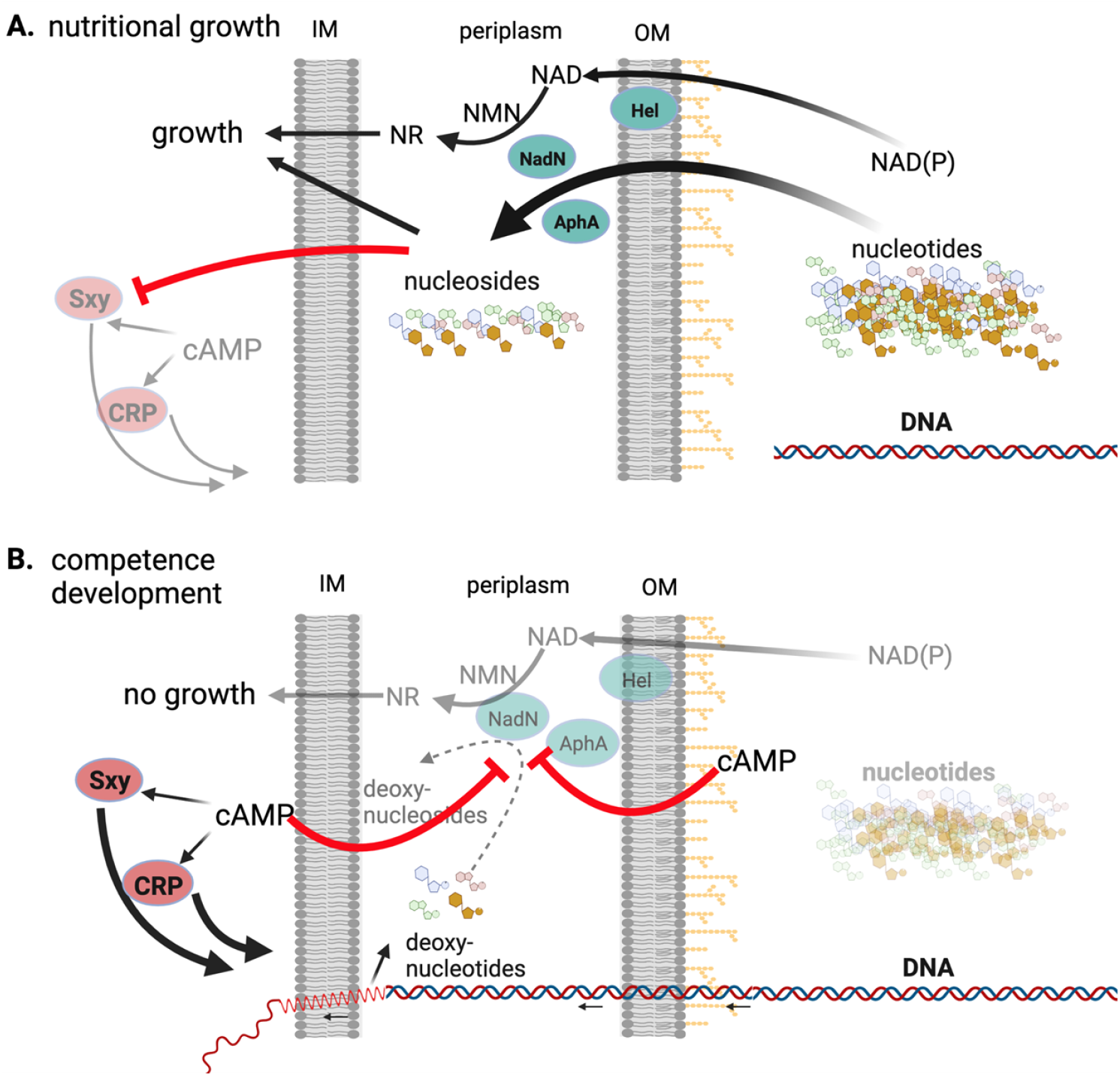
Model of the periplasmic phosphatases in coupling the cell growth with competence development in *H. influenzae* Rd KW20. (**A**) Under nutritional condition, plenty of external nucleotides enter the periplasm where they are degraded by AphA_Hi_, Hel_Hi_ and NadN_Hi_ to nucleosides, which traverse the inner membrane (IM) to the cytosol to support cell growth and inhibit the production of Sxy_Hi_, consequently the competence development. Degradation of NAD(P) to NR by Hel_Hi_ and NadN_Hi_ provides the essential growth factor V for KW20. (**B**) With exhausted carbon source, cAMP is produced by KW20. On one hand, cAMP competitively binds to AphA_Hi_, NadN_Hi_, Hel_Hi_ and inhibits their activities, slowing the generation of essential growth factors NR and pyrimidines and thus cell growth-rate. On the other hand, the generation of nucleosides is slowed by cAMP inhibiting the three enzymes, and eventually the master competence regulator Sxy_Hi_ will be produced to stimulate the gene expression required for competence development and DNA uptake. The conversion of deoxynucleotides, from degraded one DNA chain during DNA uptake, to deoxynucleosides is anticipated to be inhibited by cAMP as well. (Created with BioRender.com).

### cAMP inhibits AphA_Hi_, NadN_Hi_ and Hel_Hi_ to coordinate nutritional growth with competence development in *H. influenzae* Rd KW20

Under growth conditions with plenty of carbon sources, NAD(P), and nucleotides (Figure 6A), Hel_Hi_ cleaves NADP to NAD (11, 14), and NadN_Hi_ cleaves NAD to NMN and NR, providing the essential factor V for KW20 growth (31). AphA_Hi_ might contribute to generation of NR given its phosphatase activity. However, neither Hel_HI_ nor AphA_Hi_ is essential for KW20 growth using NAD (data not shown). Consistently, the double Δ*hel*_*Hi*_Δ*nadN*_*Hi*_ and triple Δ*hel*_*Hi*_Δ*nadN*_*Hi*_Δ*aphA*_*Hi*_ mutants showed growth patterns similar to wt strain in the sBHI medium supplemented with NR (Figure S5B). Besides NAD, all three proteins degrade various nucleotides to nucleosides (26, 31), providing both the essential pyrimidines, carbon and energy sources for cell growth.

Upon carbon starvation (Figure 6B), *H. influenzae* produces cAMP, which together with CRP_Hi_, stimulates transcription of *sxy*_*Hi*_. However, whether the *sxy*_*Hi*_ mRNA is translated to produce Sxy_Hi_ depends on exogenous nucleotides (3). Given the competitive inhibition of the phosphatase activities of Hel_Hi_, NadN_Hi_ and AphA_Hi_ by cAMP, we suggest that, under starvation, cAMP and nucleotides compete for binding to the active sites of the three enzymes (Figure 3, 4). Despite the inhibitory effect of cAMP, the excess nucleotides, especially pyrimidine nucleotides, are still gradually degraded by the three enzymes and utilized by KW20 for (slowed) cell growth (Figure 4). Meanwhile, cAMP competitively inhibits NadN_Hi_ from providing NR that is essential for growth (Figure 4), thereby gradually reducing the growth-rate. Additionally, the nucleosides produced inhibit translation of *sxy*_*Hi*_ (3). A dynamic equilibrium is reached when the exogenous nucleotides are not of a high enough concentration to compete with cAMP. This concentration threshold is expected to be lower for AMP than for GMP, given the higher affinity of AMP to AphA (Figure 2C) and the fact that cAMP cannot overcome the inhibition of competence by AMP ((3), data not shown) but by GMP (Figure 5C). Thereafter, further nucleosides are not provided and translation of the *sxy*_*Hi*_ gene is derepressed to initiate competence development. Consistently, the triple mutant was completely refractory to the inhibitory effect of AMP (Figure 5A) but the nucleoside adenosine still inhibited competence of the triple mutant (Figure 5B). Moreover, elevated levels of cAMP restored the competence inhibited by 1 mM GMP, but not so well for 1 and 5 mM guanosine (Figure 5C,5D). It is thus obvious that the competitive inhibitory effect of cAMP on the three enzymes ensures a second layer of control such that only when the external nucleotides are sufficiently depleted the competence program commences. The competitive feature is likely a key mechanism of KW20 to gauge the concentrations of cAMP and nucleotides for coordinating cell growth and competence development. As discussed, KW20 does so likely because it is auxotrophic to NAD and pyrimidine, and additionally, nucleotides provide both phosphate and carbon sources. Nucleotides are energetically expensive to synthesize from the beginning (de novo pathway); thus usage of the external nucleotides saves significant amounts of energy that may enhance ecological fitness and virulence of the organism. Consistently, in many other bacteria, such as *E. coli*, the presence of nucleotides and the nucleobases prevents the expression of genes involved in the de novo nucleotide biosynthesis via the PurR and CytR repressors (32, 33). After the extracellular nucleotide pool is sufficiently depleted and Sxy_Hi_ protein is produced, the competence regulon is fully induced, including the machinery to take up extracellular DNA (10). The dsDNA then traverses the outer membrane into the periplasm, where one strand of the dsDNA enters the cytosol, and the other strand is degraded to deoxynucleotides and released in the periplasmic space. AphA_Hi_ has similar activities towards deoxynucleotides as nucleotides (26) and, most likely, both NadN_Hi_ and Hel_Hi_ degrade deoxynucleotides as well. The accumulated deoxynucleotides potentially bind and compete for the binding site of cAMP on the three enzymes, thereby producing deoxynucleosides and phosphates. Although it is tempting to assume that these deoxynucleosides may enter the cytosol and feedback inhibit competence, it was shown before that deoxynucleosides do not inhibit competence induction of KW20 (4). Consistently, nucleotides only inhibit the early, but not the late, phase of competence induction, i.e. the translation of Sxy protein (3).

### Nucleotides regulate competence development of other bacteria

Purine nucleotides inhibit competence development of Pasteurellacean strains closely related to KW20, i.e., *Actinobacillus pleuropneumoniae* and *Actinobacillus suis (3)*. Both bacteria encode homologs of the essential competence proteins, and their competences are dependent on Sxy and CRP-cAMP. Additionally, both strains encode homologs of NadN_Hi_ (26-27% identity, 42-43% similarity), Hel_Hi_ (65% identity, 79% similarity) and AphA_Hi_ (48.4% identity, 64% similarity), except that *A. pleuropneumoniae* lacks an AphA_Hi_ homolog. Therefore, we propose that cAMP also competitively inhibit the phosphatase activities of these enzymes, to couple purine nucleotides depletion with competence development.

Nucleotides also inhibit competence of *Vibrio cholerae* (34). Natural competence in *V. cholerae* requires two signals, i.e. the presence of chitin and high cell density, and depends on cAMP, CRP and TfoX (a Sxy homolog). Recently, it was found that deletion of *cytR*_Vc_ reduced the *V. cholerae* competence to a similar level as that seen in the presence of external nucleoside cytidine (100 mM) (34). In *E. coli*, CytR represses the expression of a set of genes involved in scavenging external nucleosides (35, 36), and cytidine derepresses their expressions by direct binding to CytR_Ec_. Currently it is believed that CytR_Vc_ binds to cytidine to derepress an unknown factor that represses the expression of genes involved in natural competence development. However, besides cytidine, neither cytidine monophosphate (CMP) nor purine nucleotides were tested in the competence development of *V. cholerae* (34). Further, it is unknown whether cAMP restores the decreased competence by cytidine. *V. cholerae* encodes neither AphA_Hi_ nor Hel_Hi_ homologs, but a homolog of NadN_Hi_, i.e., UshA_Vc_ (37% identity, 52% similarity). Besides, *V. cholerae* encodes an alkaline phosphatase PhoX and CpdB_Vc_, which are homologs of PhoA_Ec_ and the cAMP specific phosphodiesterase CpdB_Ec_ protein, respectively. UshA_Vc_, PhoX, and CpdB_Vc_ are important for *V. cholerae* to use nucleotides and DNA as phosphate sources (37). Given the chemical similarity of cAMP to nucleotides, we propose that cAMP inhibits the phosphatase activities of these proteins, thus serving as a mechanism that couples competence development with nucleotides scavenging.

Another unique aspect of *V. cholerae* is that deoxycytidine also inhibits the *V. cholerae* competence (34), while deoxynucleosides do not inhibit KW20 competence (4). Therefore, it is possible that the degraded DNA chain releases deoxynucleotides in the periplasm, and that these are further degraded to deoxynucleosides that enter the cytosol and feedback inhibit the competence of *V. cholerae*. This mechanism may determine how much DNA can be taken up by *V. cholerae*.

The differential use of purine and pyrimidine nucleotides in regulating bacterial competence in KW20 and *V. cholerae* elicits some interesting questions regarding how the natural competence systems evolved. The competence systems seem to depend on both the native niches and the metabolic features of the specific bacterium. KW20 cannot synthesize NAD or pyrimidines (3). Therefore, the three proteins, NadN_Hi_, Hel_Hi_, and AphA_Hi_, likely play a house-keeping function to generate NR and pyrimidine nucleosides from the native niche of KW20, the mucus, which is known to be rich in nucleotides (38). The absolute requirement for pyrimidine and NAD for KW20 growth likely tuned the three acid phosphatases to constantly scavenging and depleting NAD and pyrimidines before the competence system is activated to take up external DNA as carbon, pyrimidine and energy sources. Despite this, it is surprising that purine but not pyrimidine suppresses competence of KW20, although KW20 has no CytR homolog but a PurR homolog. The scenario is even less clear for *V. cholerae*. The NAD, and purine and pyrimidine nucleotide synthesis pathways are complete, except for the incomplete pathway of dTTP synthesis. Thymine is therefore required for *V. cholerae* growth. *V. cholerae* produces exonucleases outside of cells to degrade DNA and release nucleotides (39, 40). Among these, dTMP may be further degraded by periplasmic phosphatases to thymidine, which enters the cytosol and is used for DNA synthesis. Since neither purine nor other pyrimidine nucleosides were tested in *V. cholerae*, the picture remains incomplete.

### Limitations of the study and perspectives

We purified a recombinant NadN_Hi_ produced in *E. coli* cytosol; however, the protein showed no phosphatase activity. This is potentially due to unique modifications in KW20, e.g. posttranslational modification, distinct signal peptide cleavage during transportation to the periplasm, oxidation etc. Given the numerous nonspecific phosphatases in the periplasm, it will be critical to quantify their concentrations under one defined condition before one can reasonably estimate the dynamic interplay between cAMP and nucleotides, nucleosides, and their effect on competence development. For *V. cholerae*, it remains interesting to test if purine nucleotides also suppress competence development and if cAMP restores it. Although it is anticipated so, KW20 and *V. cholerae* have distinct metabolic features discussed above that somehow may contribute to the differential effects of purine and pyrimidine in both organisms.

### Materials and Methods

#### Bacterial strains, growth, media and antibiotics

The strains and primers used in this project were listed in the Table S1 and S2, respectively. Antibiotics used for the specific strains are listed in Table S1. The *E. coli* K-12 MG1655 is the wild type strain used. Lysogeny broth (LB, containing 10 g tryptone (Oxoid), 5 g yeast extract (Oxoid), and 10 g NaCl (SIGMA) per liter of distilled water) was the primary medium used for *E. coli* growth. For competence assays *H. influenzae* Rd KW20 was the wild type used. A brain heart infusion medium (BHI, Becton and Dickinson) supplemented with 15.4 μM hemin and 14.6 μM nicotinamide adenine dinucleotide (NAD) (hereby referred to as sBHI) was the rich medium for this strain. For competence induction, M-IV medium was prepared as described before (41). LB and sBHI agar plates contained 1.5% agar (Difco).

#### Competence assays

Natural competence in *H. influenzae* cells was performed as in (3). Briefly, competence was induced by transferring early log-phase cells (OD_600_≈0.2) grown in sBHI to the M-IV starvation medium, and incubated for 1 hour at 100 rpm, 37°C. If cAMP was included in the assay, it was added at the onset of competence induction. When testing for the repressive effect of nucleotide precursors on competence development, this compound was added 15 min into competence induction. Cells were next incubated with 1 μg/ml purified chromosomal *H. influenzae* DNA encoding novobiocin resistance (from YZ1080) for 1 hour and were then serial diluted in BHI and spotted on sBHI plates with or without 2.5 μg/ml novobiocin (SIGMA). Competence development was measured by dividing the novobiocin resistant CFU by the total CFU. Note that, the YZ1080 strain was transformed with the MAP7 DNA (41) containing seven antibiotic markers in KW20 genomic DNA and selected on sBHI supplemented with 2.5 μg/ml novobiocin.

#### Deletion of aphA_Hi_, hel_Hi_ and nadN_Hi_ in H. influenzae Rd KW20

For deletion of *aphA*_*Hi*_, *hel*_*Hi*_ and *nadN*_*Hi*_, the natural competence of *H. influenzae* was utilized to replace each of these genes with an antibiotic marker. Each of the antibiotic marker genes are proceeded with the promoter sequence (TAAATTGAACTTTTTTCTTCATCAGAACTCAAAAACAACGTTCTCTGCCTAATTGAATTGGGCAGAGAAAATATTAAACCCATCATTTAATTAAGGATATTTATCAA) from an constitutively expressed KW20 gene *omp26*. Then, ca. 1000 bp homologous DNA sequences upstream and downstream of the *aphA*_*Hi*_, *hel*_*Hi*_, *nadN*_*Hi*_ genes were fused to the antibiotic marker via overlap PCR, to facilitate the recombination between the endogenous genes and the antibiotic marker sequences. As an example, for deletion of *aphA*_*Hi*_, primers DaphA-1 (*AACGGCGCGCAATTTCAGTTTTACC*) and DaphA-2 (CTGATGAAGAAAAAAGTTCAATTTA*tgcctttcctcacaaacgctgatt*) were used to PCR amplify the upstream sequence of *aphA* gene, and primers DaphA-3 and DaphA-4 were used to amplify the *omp26* promoter DNA, and primers DaphA-5 and DaphA-6 were used to amplify the chloramphenicol resistant marker for pWRG99 plasmid ref, and primers DaphA-7 and DaphA-8 were used to amplify the downstream sequence of *aphA*_*Hi*_ gene. The four DNA sequences were then fused together via overlap PCR to obtain the DNA to be fed to the competent KW20 cells and selected on sBHI media supplemented with chloramphenicol 4 µg/ml. Similarly, for deletion of *hel*_*Hi*_, primers pYZ653 and pYZ654, pYZ655 and pYZ656, pYZ657 and pYZ658, and pYZ659 and pYZ660 were used to amplify the corresponding four DNA fragments that were fused together via overlap PCR. The same approach was done for deletion of *nadN*_*Hi*_ but using primers pYZ661 and pYZ662, pYZ655 and pYZ656, pYZ663 and pYZ664, and pYZ665 and pYZ666. The final DNA constructs were then added to competent *H. influenzae* cells before these were plated onto selective plates (kanamycin 5 µg/ml for *hel*_*Hi*_ deletion and spectinomycin 20 µg/ml for *nadN*_*Hi*_ deletion). To make the double and triple mutants of *aphA*_*Hi*_, *hel*_*Hi*_ and *nadN*_*Hi*_, the chromosomal DNA of the single deletion strains of *hel*_*Hi*_ and *nadN*_*Hi*_ was used to transform the wt or *aphA*_*Hi*_ single deletion strains. The mutants were selected on sBHI agar plate supplemented with appropriate antibiotics. Besides, nicotinamide riboside was required in sBHI to support the growth of mutants with *nadN*_*Hi*_ deleted. Positive transformants were first identified via colony PCR and then confirmed by sequencing (using primers pYZ688 and pYZ689 for *hel*_*Hi*_ deletion and pYZ690 and pYZ691 for *nadN*_*Hi*_ deletion).

#### Plasmid constructions

pET28a-*aphA*_*Ec*_-his

The primers pYZ106 and pYZ107 were used to amplify the *aphA*_*Ec*_ and constructed into the plasmid vector pET28a via the NcoI and NdeI restriction sites.

pET28a-his-*aphA*_*Ec*_

The primers pYZ194 and pYZ195 were used to amplify the *aphA*_*Ec*_ and constructed into the plasmid vector pET28a via the NcoI and NdeI restriction sites.

pET24d-his-sumo-*aphA*_*Ec*_

The primers pYZ368 and pYZ369 were used to amplify the *aphA*_*Ec*_ and constructed into the plasmid vector pET24d (Novagene) via the BamHI and HindIII restriction sites. Then, the linker region between *aphA* and sumo was removed via quickchange mutagenesis by using primers pYZ370 and pYZ371.

pET28a-his-*aphA*_*Hi*_

The primers pYZ198 and pYZ199 were used to amplify the *aphA*_*Hi*_ and constructed into the plasmid vector pET28a via the NcoI and NdeI restriction sites.

pET28a-his-*hel*_*Hi*_

The primers pYZ200 and pYZ201 were used to amplify the *hel*_*Hi*_ and constructed into the plasmid vector pET28a via the NcoI and NdeI restriction sites.

pET24d-his-sumo-*hel*_*Hi*_

The primers pYZ372 and pYZ373 were used to amplify the *hel*_*Ec*_ and constructed into the plasmid vector pET24d via the BamHI and EcoRI restriction sites. Then, the linker region between hel and sumo was removed via quick-change mutagenesis by using primers pYZ374 and pYZ375.

pET28a-his-*nadN*_*Hi*_

The primers pYZ790 and pYZ791 were used to amplify the *nadN*_*Hi*_ sequence with the signal peptide and constructed into the plasmid vector pET28a via the BamHI and NdeI restriction sites.

#### Protein purification

*E. coli* strains were inoculated into LB medium supplemented with the appropriate antibiotic and incubated ON at 37°C shaking 160 rpm. The next morning, the culture was diluted back into LB-medium (1:500 dilution) and grown at 37°C, 160 rpm to OD_600_=0.6-0.8, before the culture was induced with 0.5-1 mM IPTG and incubated for 4 hours at 37°C, 160 rpm. Cells were harvested by centrifugation (5000 rpm, 10 min, 4°C), resuspended in cold PBS and pelleted (4000 rpm, 10 min, 4°C). The pellet was resuspended in cold lysis buffer (5% glycerol 50 mM Tris pH=7.6, 150 mM NaCl, 10 mM imidazole) supplemented with β-mercaptoethanol (0.35 μl/ml sample) and one tablet of cOmplete^TM^ Protease Inhibitor Cocktail and was then lysed in a Branson sonicator for 8 min (2 min ON/4 min OFF) with an amplitude of 60%. The lysate was centrifugated (14000 rpm, 40 min, 4°C) and the supernatant was loaded onto Ni-NTA resins on a Poly-Prep column and allowed to drip through via gravity. The resins were then washed in cold wash buffer (5% glycerol 50 mM Tris pH=7.6, 150 mM NaCl, 20 mM imidazole), eluted in 700 μl elution buffer (5% glycerol 50 mM Tris pH=7.6, 150 mM NaCl, 500 mM imidazole) and further purified via the ÄKTA system on a Superdex 200 Increase 10/300 GL column.

#### DRaCALA screening and assays

The ^32^P labeled cAMP used for the DRaCALA were produced as described previously (17). The screening of the cAMP binding proteins from the ASKA *E. coli* ORFeome library was performed essential the same as in (17, 20). For determining the dissociation constant of cAMP binding to AphA, purified AphA protein was mixed with ^32^P-cAMP in a binding buffer composed of 40 mM Tris (pH 7.5), 100 mM NaCl, 10 mM MgCl_2_). This mixture was next serial diluted in the same binding buffer containing ^32^P-cAMP (2 nM), and the samples were then incubated at room temperature for 5 min before they were spotted on a nitrocellulose membrane and visualized in a Typhoon FLA-7000 phosphorimager. For DRaCALA competition assays, the cold cAMP, AMP or GMP at varied concentrations were individually incubated with AphA and ^32^P-cAMP (2 nM) in the same binding buffer as above. Samples were then spotted and visualized as described above. The fraction of bound were quantified as described previously (17).

#### Electrophoretic Mobility Shift Assay

A piece of *H. influenzae* chromosomal DNA (200 bp) containing the uptake sequence UPS ACCGCACTT was PCR amplified using primers pYZ609/pYZ610, which are labelled with cy5. MAP7 DNA (41) was used as template. The DNA was subsequently purified from an agarose gel using Monarch® Gel Extraction Kit. For the EMSA, purified AphA_Hi_ protein was incubated with this DNA in a buffer composed of 20 mM Tris-HCl (pH 7.5), 100 mM KCl, 2 mM MgCl_2_, 1 mM β-mercaptoethanol, 50 μg/ml BSA and 0.1 mg/ml salmon sperm DNA in a total volume of 20 μl. Following 20 min of incubation at 37°C, 3 μl 80% glycerol was mixed into the samples that were next loaded into a 5% Mini-PROTEAN® gel (Bio-Rad). The gel was run at 40 V for 10 min, then at 80 V for 1 h and was finally visualized in an Image Quant LAS4000 scanner.

#### Biochemical assays

To assess the enzymatic reaction of AphA, kinetic assays were performed using pPNN as the substrate as in (26). Freshly purified AphA protein was resuspended in a buffer consisting of 50 mM sodium acetate (pH 5.6) and 0.01% TritonX-100 and was subsequently transferred into a reaction buffer composed of 50 mM sodium acetate (pH 5.6), 0.1 M NaCl, 1 mM MgCl_2_, 0.01% Triton X-100 and varying concentrations of cAMP. The enzymatic reaction was started by adding varying concentrations pPNN to the reaction mixture and was terminated by transferring 60 μl of the reaction mixture into 100 μl 3 M NaOH in a clear Greiner flat bottom 96 well plate at defined timepoints. The amount of reaction product generated from pNPP was quantitated in a plate reader at 405 nm.

---

**Table S1:** Bacterial strains used in this study

| Strain | Relevant features | Reference |
| --- | --- | --- |
| YZ9 | DH5α pET28a, Kan | Novagene |
| YZ576 | pET24d; Kan | Novagene |
| YZ16 | BL21 DE3 | Laboratory stock |
| YZ142 | MG1655, wild type *E. coli* | Laboratory stock |
| YZ308 | pCA530-Ulp1-His, Kan | Laboratory stock |
| YZ272 | Rd KW20, wild type *H. influenzae* | Laboratory stock |
| YZ97 | Bl21 DE3 pET28a-*aphA*-his | This study |
| YZ253 | Bl21 DE3 pET28a-his-*aphA*<sub>Ec</sub>, Kan | This study |
| YZ577 | Bl21 DE3 pET28a-his-sumo-*aphA*<sub>Ec</sub>, Kan | This study |
| YZ254 | Bl21 DE3 pET28a-his-*aphA*<sub>Hi</sub>, Kan | This study |
| YZ255 | Bl21 DE3 pET28a-his-*hel*<sub>Hi</sub>, Kan | This study |
| YZ579 | Bl21 DE3 pET28a-his-sumo-*hel*<sub>Hi</sub>, Kan | This study |
| YZ1340 | Bl21 DE3 pET28a-his-*nadN*<sub>Hi</sub>, Kan | This study |
| YZ278 | Rd KW20 *aphA::cam*, Cam | This study |
| YZ991 | Rd KW20 Δ*hel::kan*, Kan | This study |
| YZ993 | Rd KW20 Δ*aphA::cam*; Δ*hel::kan*, Cam, Kan | This study |
| YZ995 | Rd KW20 Δ*hel::kan*; Δ*nadN::spec*, Kan, Spec | This study |
| YZ997 | Rd KW20 Δ*aphA::cam*; Δ*hel::kan*; Δ*nadN::spec*, Cam, Kan, Spec | This study |
| YZ1080 | Rd KW20, Nov | This study |

**Table S2:**
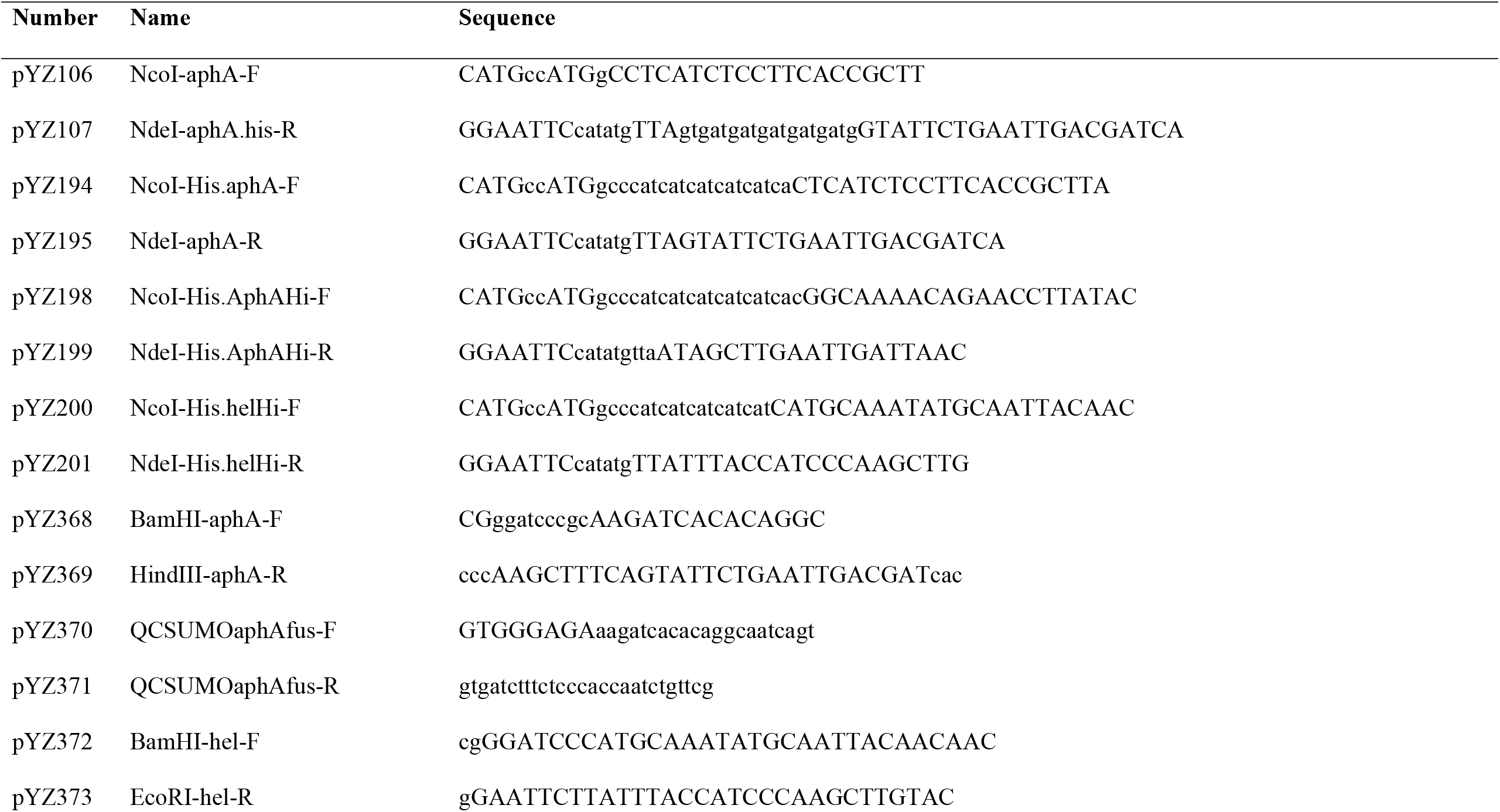
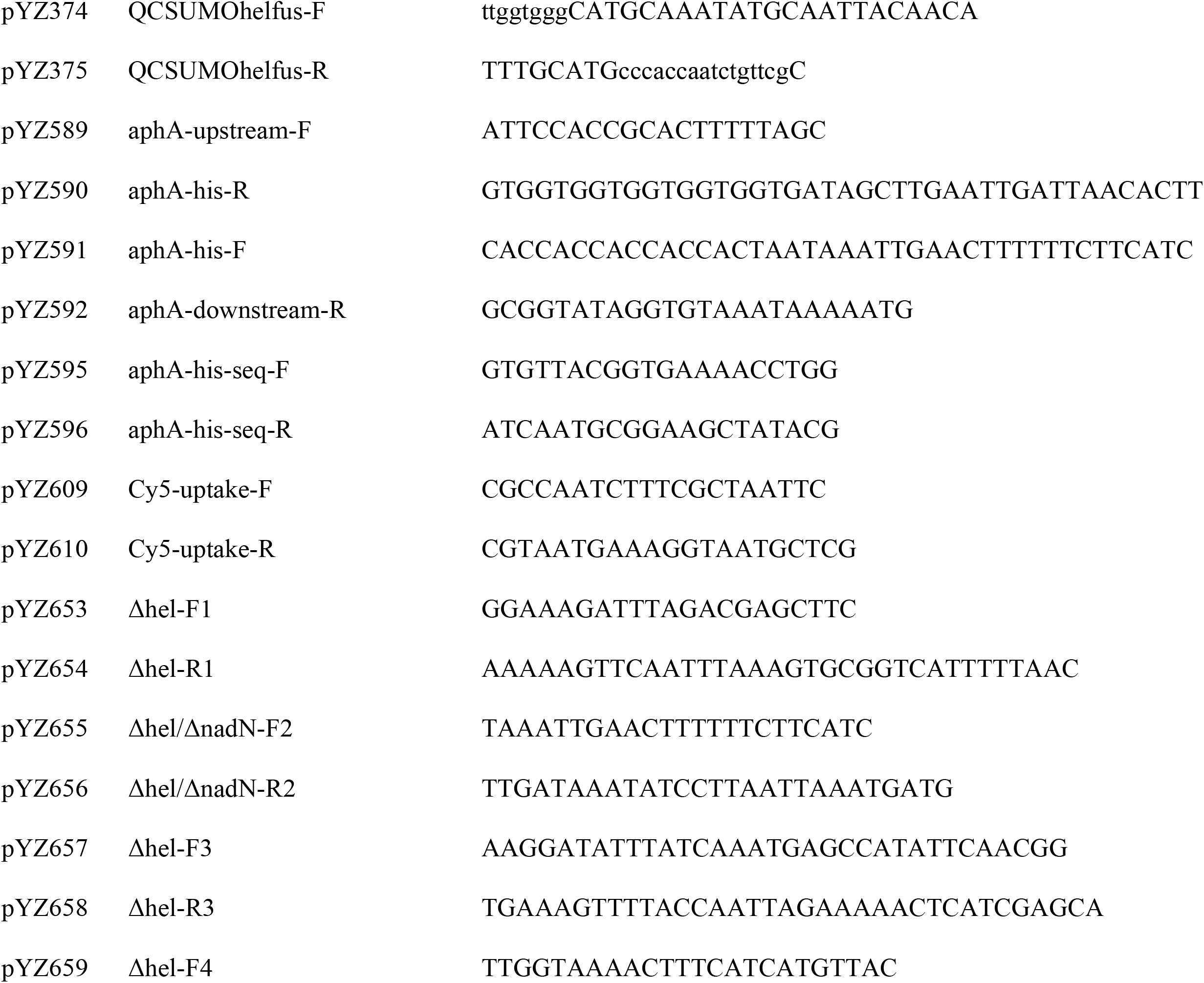
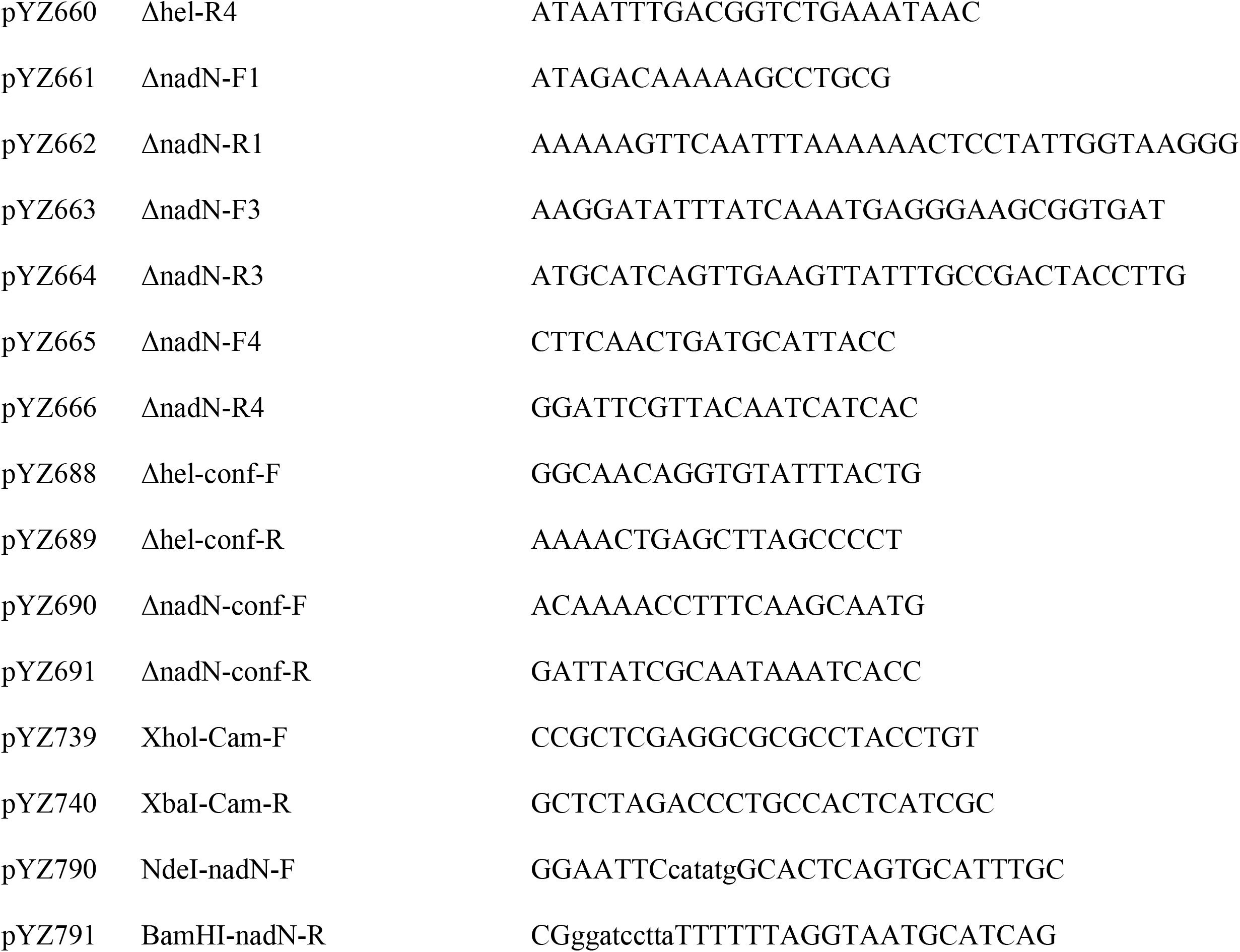
Primers used in this study

## Acknowledgement

We are grateful to Kenn Gerdes for the early-phase support of this project and critical reading of the manuscript, and to Rosemary Redfield for sharing the MAP7 DNA and useful discussions. We wish to express our thanks to Farshid Jalalvand for making the KW20 *Δaph*A_*Hi*_::cat mutant. This study was supported by a Novo Nordisk Foundation

Project Grant (NNF19OC0058331) to Y.E.Z.

---

**Figure S1.**
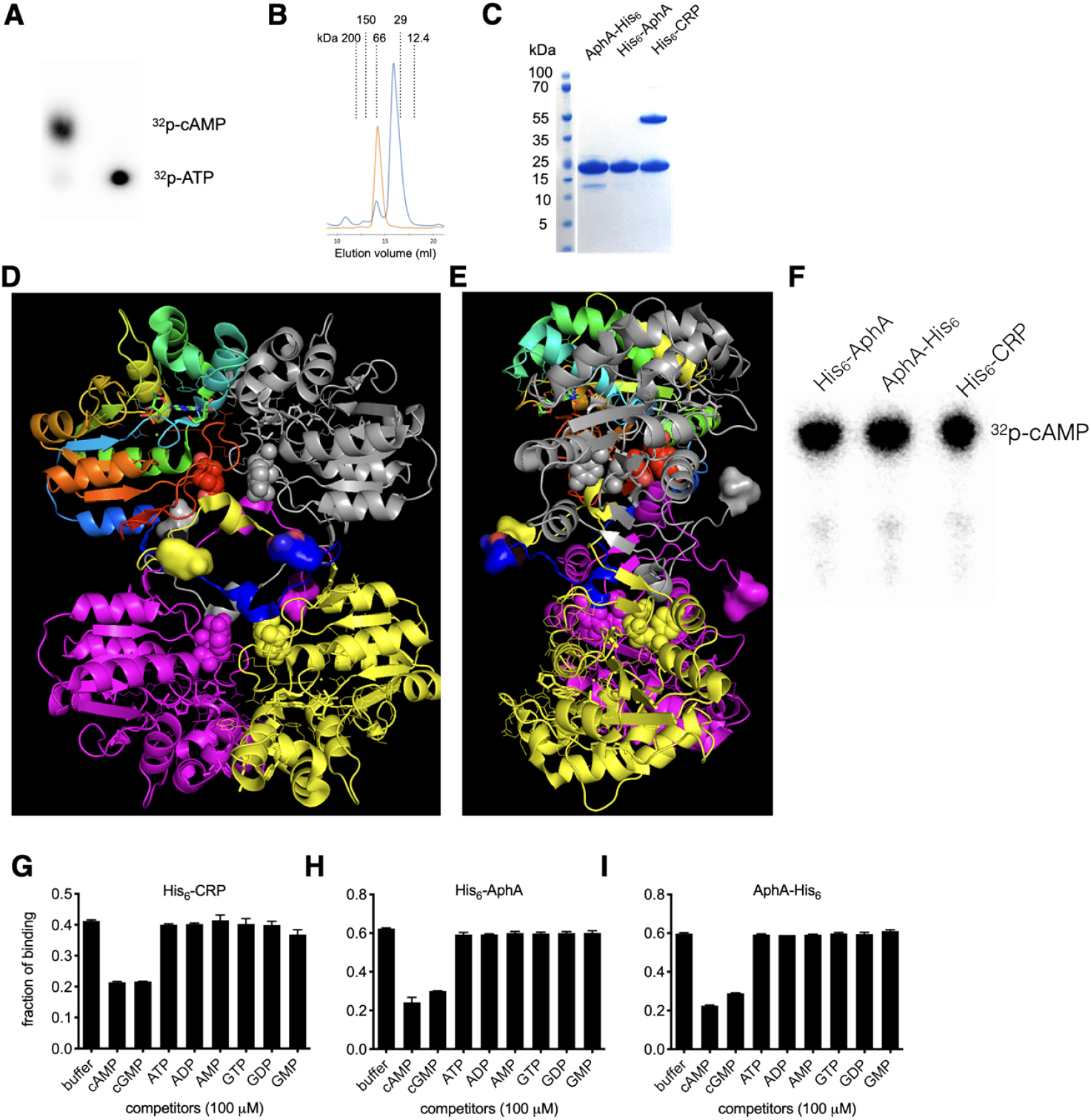
(**A**) Thin Layer Chromatography (TLC) plate showing the synthesis of ^32^p-labelled cyclic AMP [p^32^-α-cAMP] from p^32^-α-ATP. (**B**) The FPLC profiles of N-(orange) and C-(blue) 6-histidine tagged AphA_Ec_. The sizes of the standard proteins are marked on the top. (**C**) An SDS-PAGE gel picture of purified His_6_-AphA_Ec_, AphA_Ec_-His_6_, and His_6_-CRP_Ec_ proteins. (**D**) Structure of *E. coli* AphA_Ec_ (2B82) presented in cartoon model with the N- and C-termini shown in surface and sphere models, respectively. The four monomers were colored differently. (**E**) Side view of the structure of AphA_Ec_ (2B82). (**F**) TLC plate showing that the p^32^-α-cAMP was not degraded by AphA_Ec_ (10 µM) after 20 min at 37°C. (**G**,**H**,**I**) Quantitation of DRaCALA competition assay by using purified His_6_-AphA_Ec_, AphA_Ec_-His_6_, and His_6_-CRP_Ec_, with the presence of buffer, cold cAMP, cGMP, ATP, ADP, AMP and GTP, GDP, GMP (each at 100 µM). Two biological replicates were performed, and the average and standard deviation of mean are shown.

**Figure S2.**
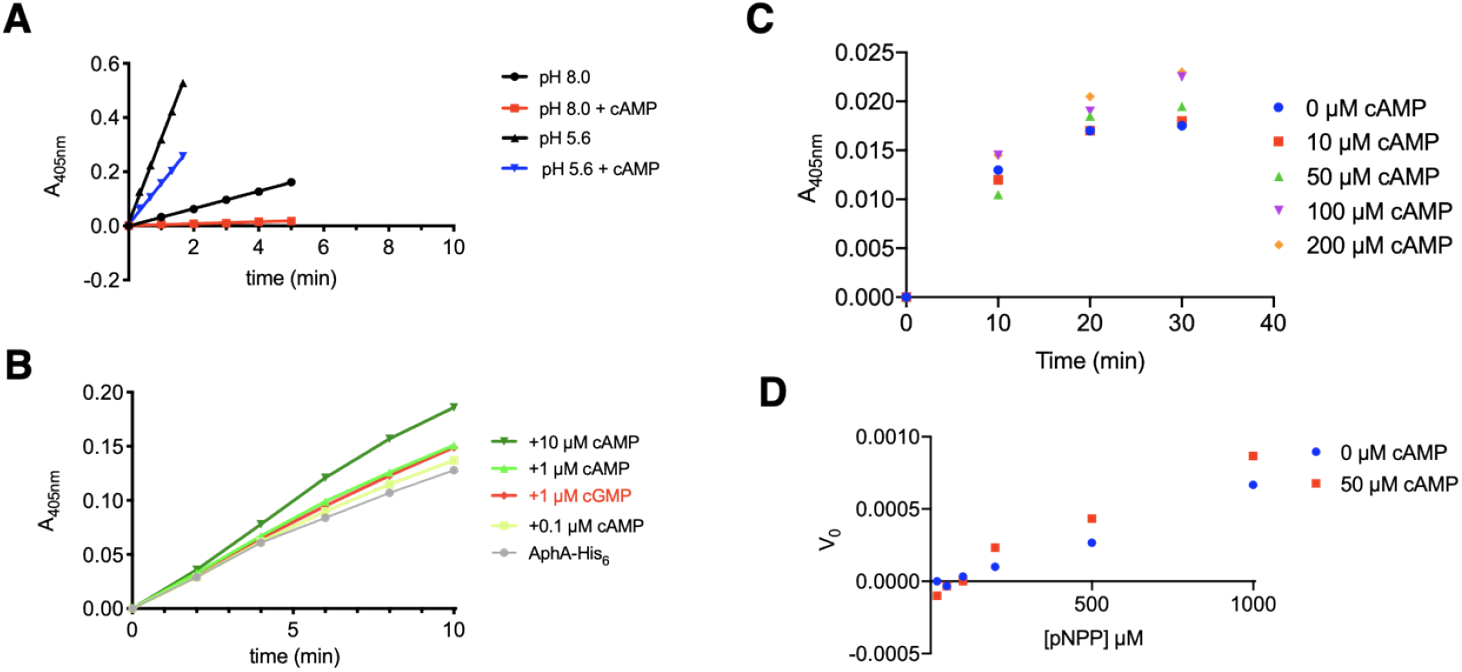
cAMP inhibits the catalytic activity of His_6_-AphA_Ec_ (**A**) but stimulates AphA_Ec_-His_6_ (**B**). The acid phosphatase assay was performed by using pNPP (p-Nitrophenyl Phosphate, 1 mM) as substrate and 15 nM of each AphA_Ec_ protein at pH 8.0 and 5.6 (see text and Method for details). cAMP (and cGMP) at various concentrations were added to assess their effects on the AphA_Ec_ activity. (**C**) Time course of AphA_Ec_-His_6_ cleaving pNPP in the absence and presence of cAMP. (**D**) Michaelis-Menten curves of AphA_Ec_-His_6_ cleaving pNPP in the absence and presence of 50 µM cAMP.

**Figure S3.**
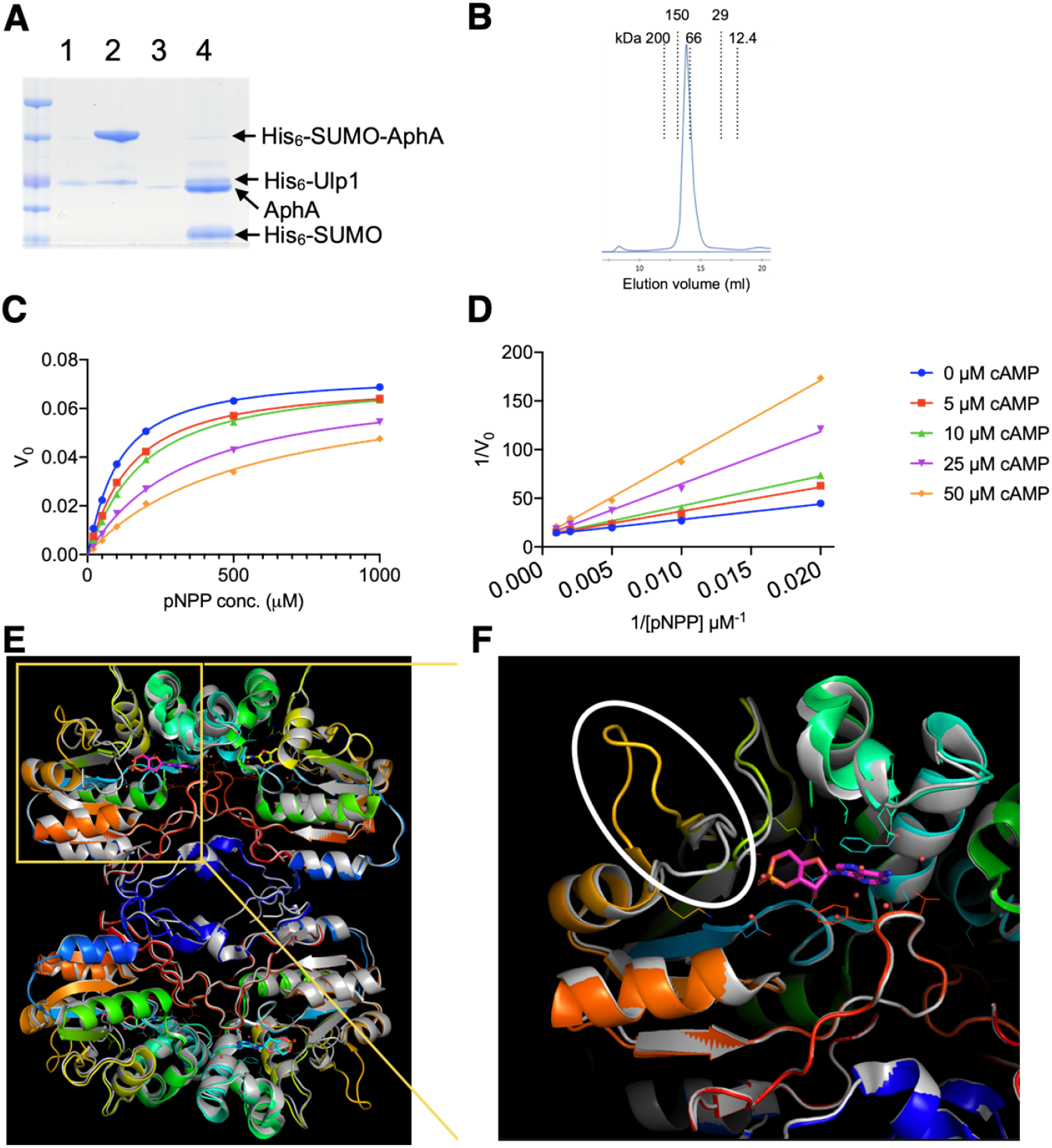
(**A**) An SDS-PAGE gel test of non-cleaved and cleaved His_6_-SUMO-AphA_Ec_ protein. 1, His_6_-Ulp1; 2, His_6_-SUMO-AphA_Ec_; 3, tag-less AphA_Ec_ after cleavage; 4, the cleavage reaction mixture. (**B**) FPLC profile of the cleaved tag-less AphA_Ec_. (**C**) Michaelis-Menten curves and (**D**) the Lineweaver–Burk plot of His_6_-AphA_Ec_ cleaving pNPP in the absence or presence of cAMP. (**E**) Superposition of *E. coli* AphA_Ec_ structure (2B82, grey) to the *Salmonella typhimurium* AphA in complex with cAMP (1U5Z, color). (**F**) Zoom-in of the cAMP binding pockets in 1U5Z (E) and the superimposed 2B82. The main conformational difference between the two structures is circled.

**Figure S4.**
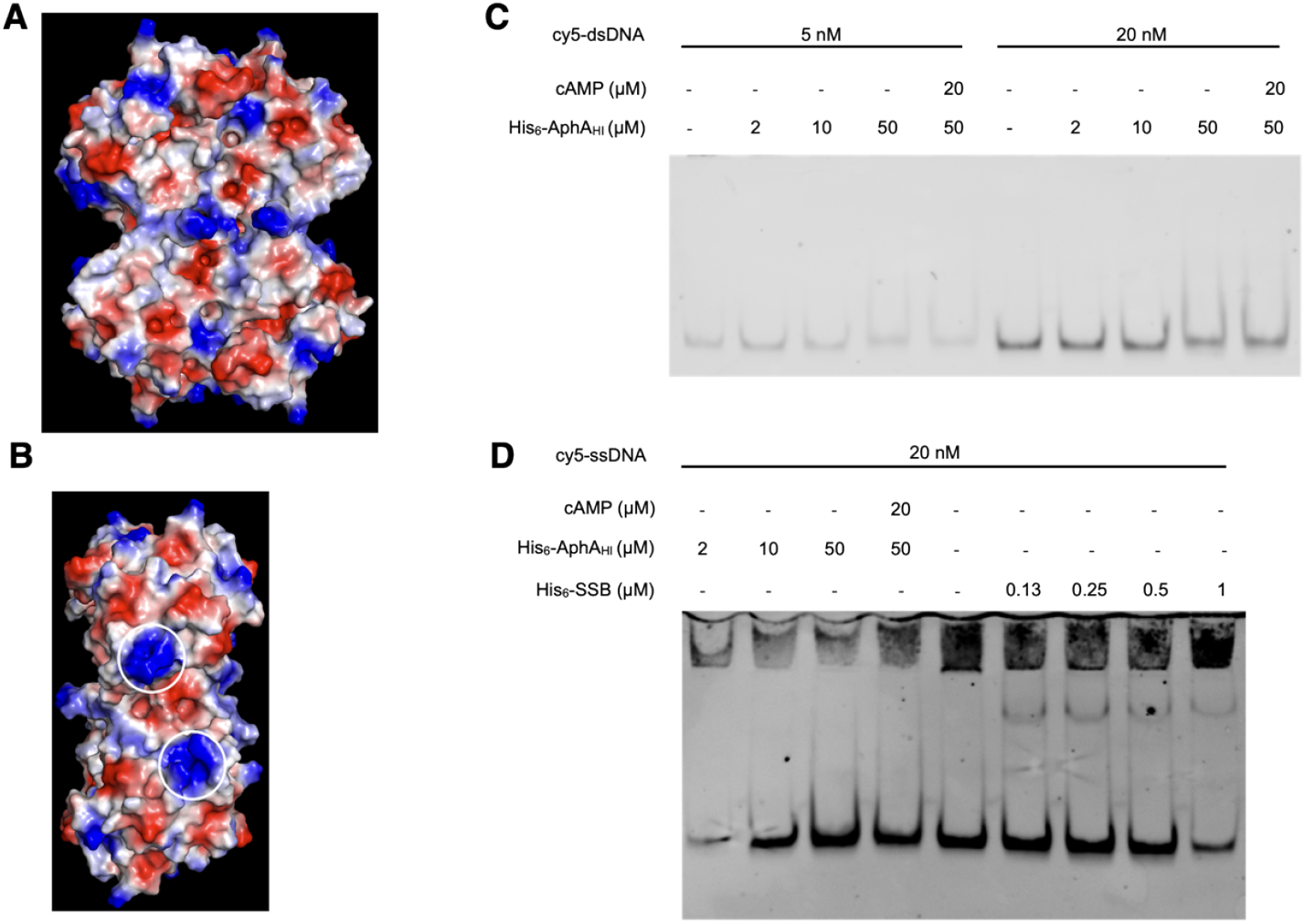
The front (**A**) and side (**B**) views of the surface charge of *E. coli* AphA_Ec_ (PDB 2B82). Blue and red indicate positively and negatively charged area, respectively. (**C**) Gel retardation assay testing the binding of His_6_-AphA_Hi_ (µM) to a 200-bp long double strand dsDNA (5 and 20 nM) amplified from *H. influenzae* chromosome, which was labelled with cy5 at the 5’-ends. (**D**) Gel retardation assay similar as in panel C, except that the dsDNA was denatured to single stranded ssDNA. The *E. coli* single-strand DNA binding protein (His_6_-SSB) was used as positive control.

**Figure S5.**
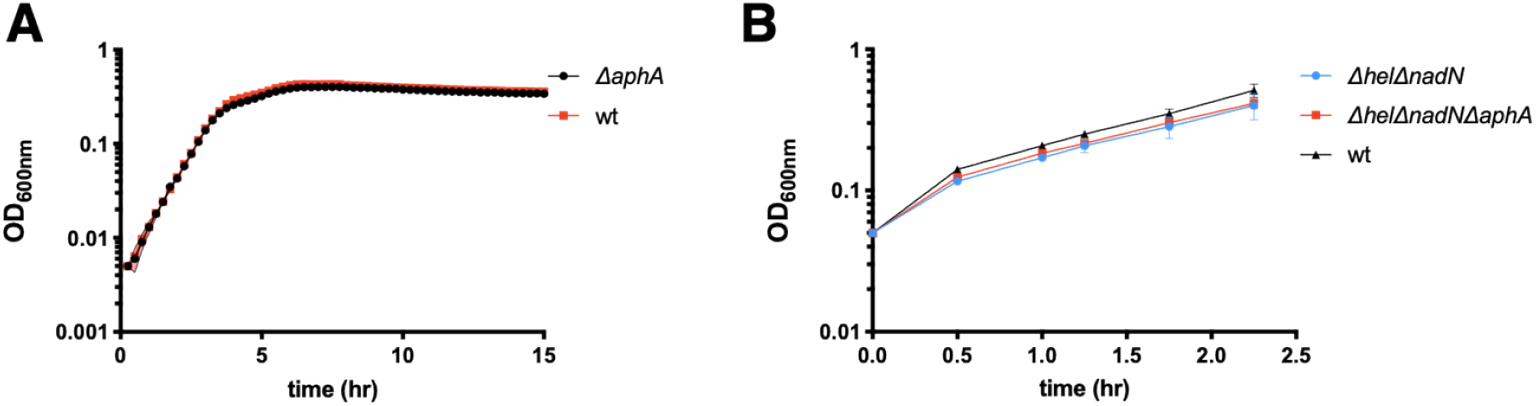
Growth curves of various strains of *H. influenzae* Rd KW20 in sBHI media supplemented with NAD (**A**) or NR (nicotinamide riboside) (**B**). Two biological replicates were performed, and the average and standard deviation (SD) are shown.

**Figure S6.**
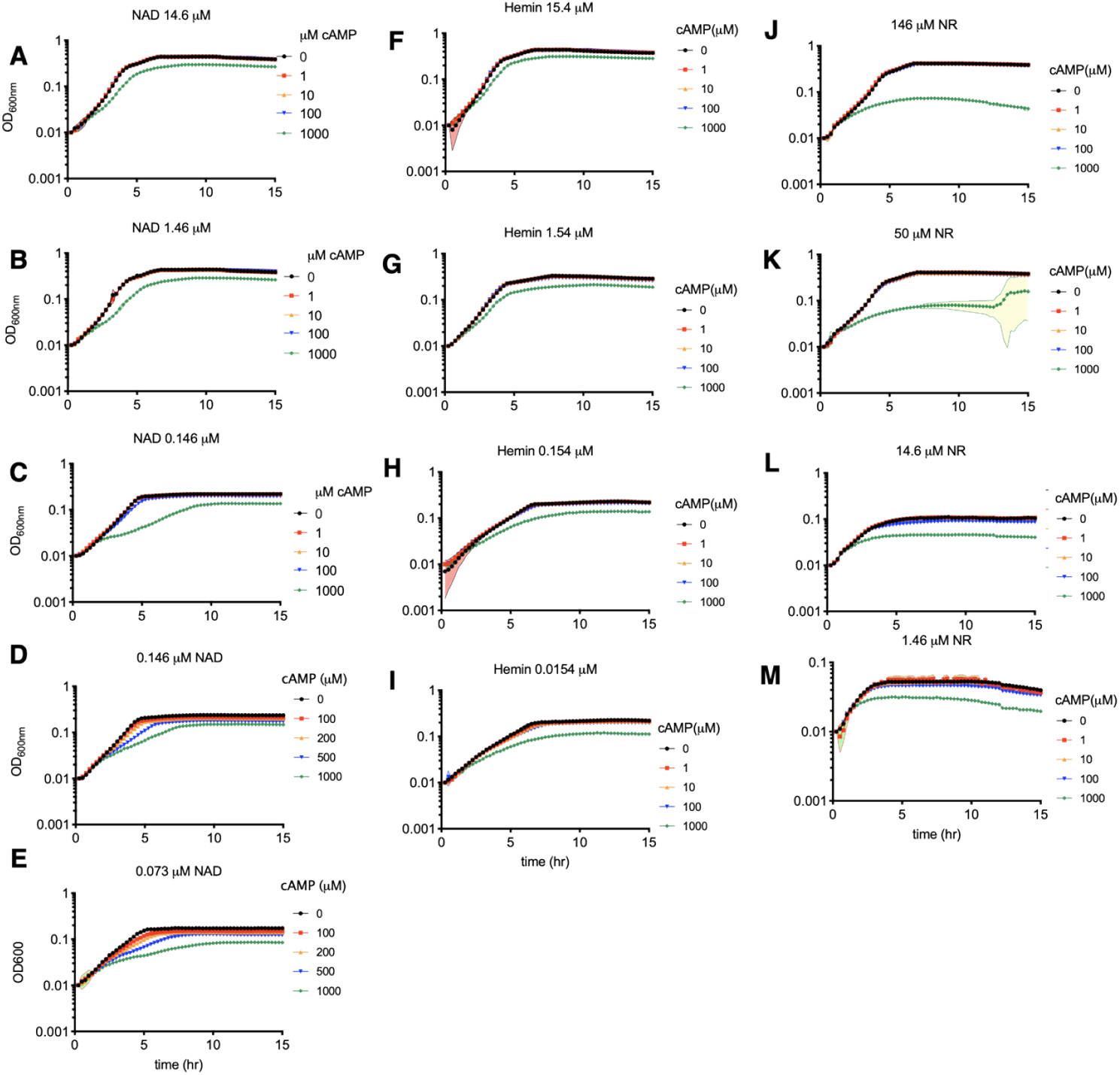
Growth curves of the *H. influenzae* Rd KW20 in sBHI medium supplemented with varied concentrations of NAD (**A-E**), hemin (**F-I**), and NR (nicotinamide riboside) (**J-M**) in the presence of differential concentrations of cAMP. Three biological replicates were performed, and the average and standard deviation of mean are shown.

